# Machine learning informs mitigation strategies for nitrous oxide emissions from wastewater operations

**DOI:** 10.1101/2025.08.31.673305

**Authors:** Gnanaraj Augustine, Kartik Chandran

**Affiliations:** Department of Environmental Engineering, Columbia University, New York, 10027, USA

**Keywords:** activated sludge, biological nutrient removal, greenhouse gas, machine learning, mitigation, nitrous oxide, performance

## Abstract

This study focused on the development of machine-learning-(ML) based strategies for mitigating nitrous oxide (N_2_O) emissions from various wastewater treatment systems in the United States measured using a benchmark USEPA-endorsed protocol. Results revealed that in general, process performance correlated with higher N_2_O emissions. Among the models evaluated, the Extreme Gradient Boosting (XGBoost) model demonstrated strong predictive performance for aerobic (R^2^=0.97), and anoxic (R^2^=0.87) zone process data. Specifically, the unsupervised principal component analysis (PCA), along with the trained ML models, demonstrated that local variables including zone-specific dissolved oxygen (DO), ammonia (NH_3_), and nitrite (NO_2_^-^) concentrations and global variables including effluent nitrite and nitrate as key contributors towards N_2_O emissions from both aerobic and anoxic zones of the process bioreactors. The operating regions associated with lower predicted N_2_O emissions included operations of aerobic and anoxic zones at DO < 4 mg O_2_ L^-1^ and < 1 mg O_2_ L^-1^ respectively, coupled with appropriate solids retention times (SRTs) that maximize process performance. The scenario analysis using the trained ML model predicted 7.8-fold (95% CI: 2.2 to 52.8-fold) and 8.8-fold (95% CI: 3.83 to 21.24-fold) higher N_2_O flux at low DO/high NH_3_ and high DO/high NO_2_^-^ conditions in aerobic and anoxic zones, respectively. This model predicted responses were consistent with the established nitrifier denitrification and N_2_O reductase inhibition pathways. Accordingly, our results underscore the utility of ML models interpreted in conjunction with bioprocess fundamentals for predicting and mitigating N_2_O emissions, while concomitantly improving wastewater treatment operations.

## 1. Introduction

Nitrous oxide (N_2_O) is a potent greenhouse gas (GHG) and ozone depleting substance^1,2^. Wastewater treatment plants (WWTPs) have been increasingly recognized as substantial anthropogenic sources of N_2_O emissions^3–7^. On one hand, increasingly stringent effluent nitrogen limits from wastewater treatment plants have spurred a global effort to develop and implement engineered biological nutrient removal (BNR) processes to improve water quality^4^. On the other hand, suboptimal operating conditions and process configurations associated with BNR operations can exacerbate N_2_O production^8^.

During typical WWTP nitrification operations, chemolithoautotrophic ammonia-oxidizing bacteria (AOB) and nitrite-oxidizing bacteria (NOB) sequentially oxidize ammonia (NH_3_) to nitrite (NO_2_^-^) and nitrate (NO_3_^-^), respectively, using oxygen as the preferred terminal electron acceptor^9^. During denitrification, NO_3_^-^ is reduced to dinitrogen (N_2_) gas, coupled with the utilization of organic or inorganic electron donors^9^. In nitrifying WWTPs, AOB produce nitric oxide (NO) and N_2_O via nitrifier denitrification under limiting dissolved oxygen (DO) and non-limiting NO_2_^-^ concentrations^6,10–15^. Also, under non-limiting DO and NH_3_ concentrations, AOB produce N_2_O during the oxidation of hydroxylamine (NH_2_OH)^10,16^. N_2_O can also be produced through chemical oxidation of NH_2_OH^10,16^. During typical engineered denitrification, chemoorganoheterotrophic bacteria produce N_2_O under operating conditions that favor inhibition of nitrous oxide reductase (Nos) ^16^, for instance, associated with inhibition by oxygen and NO_2_^-12,17,18^. Despite the contribution of both nitrification and denitrification in N_2_O production, it has now been established that both *production* and *emission* of N_2_O from aerated zones are higher than from non-aerated zones at WWTPs^4^.

The characterization and minimization of N_2_O emissions from WWTPs has assumed widespread and critical importance as wastewater utilities aim to minimize the overall carbon footprint of treatment operations, while adhering to increasingly stringent discharge regulations^19^. Previous N_2_O footprinting studies performed at full-scale BNR and non-BNR WWTPs around the United States reported N_2_O emission factors (EFs) in the range 0.01-1.8% relative to the influent TKN load^4^. Other full-scale studies have reported widely varying EFs in the range 0.001-25.4% of the influent load^4,20^.

N_2_O production is directly correlated with nitrogen conversion dynamics in WWTPs, further indicating the potential of N_2_O emissions as BNR performance indicators^4–6,20^. Various operational and environmental factors such as solids retention time (SRT), specific AOB activity, temperature, and influent loading patterns, contribute to both N_2_O emissions and overall process performance in BNR systems^4–7,20^. BNR operations at sub-optimal conditions, such as excessively high or low DO concentrations combined with inadequate SRT, can lead to transient or permanent NH_3_ or NO_2_^-^ accumulation. Such accumulation is symptomatic of inadequate nitrification and denitrification performance and can be correlated, in turn, to NO and N_2_O production and emission^5–10^. Mechanistic models have been used to predict and reduce N_2_O emissions in the BNR systems from WWTPs, ranging from models based on the activated sludge framework^21^ to models based on gene- and protein-expression^15,22–25^. Other data-driven models have also been proposed, which when trained on robust and representative datasets, offer a balance between the mathematical and metabolic intricacies relating to N_2_O production and their practical end-application for process control^4,26^. Data-driven models such as unsupervised Principal Component Analysis (PCA) and supervised machine learning (ML) models have shown greater insights into improving process performance and identifying factors contributing to N_2_O emissions^27–32^. Supervised ML models have demonstrated the potential for predicting N_2_O emissions from full-scale BNR operations, and studies have demonstrated the use of ML models focused on predicting effluent water quality^32–36^ or primarily N_2_O emissions.^37–40^, using different models and approaches such as hyperparameter optimization (HPO)^40^.

In this study, we subjected both historical and currently obtained datasets to unsupervised PCA and supervised ML modeling approaches to uncover the complex interplay between operational strategies, N_2_O emissions and effluent water quality, and to develop actionable insights into WWTP operating strategies that minimize both *gaseous* and *aqueous* nitrogenous discharges. Through this research, we offer new insights into concurrently maximizing BNR process efficiency and achieving decarbonization of BNR operations, thereby promoting a more sustainable wastewater sector.

## 2. Materials and Methods

### 2.1. Field Data Collection using a USEPA Approved Protocol to Measure N_2_O Emissions from BNR Processes

We performed N_2_O measurement campaigns (2023-2024) at three different BNR wastewater treatment plants, representing five distinct mainstream and sidestream process configurations located in the Northeastern and Mid-Atlantic regions of the United States. For each process configuration, N_2_O measurements were collected continuously over 24-hour monitoring campaigns to capture diurnal variability. The mainstream four-pass step-feed BNR was monitored for 50 days at low temperature (18±3°C) and 52 days at high temperature (23±3°C). The partial denitratation anammox (PdNA) process was monitored for 8 days at low temperature (18±1.7°C) and 7 days at high temperature (25±2.8°C), respectively. The ammonia-based aeration control (ABAC) process was monitored for 2 days at 26±2.2°C, whereas the fermentate-fed pilot denitrifying system was monitored for 1 day at 25±0.7°C. The sidestream SHARON process was monitored for over 47 days under elevated influent temperature conditions (**Table 1**). Repeated daily measurements within each configuration represented temporal observations of a single treatment system and were therefore treated as repeated monitoring events rather than independent replicates. Accordingly, inter-configuration comparisons are presented as descriptive summaries rather than inferential statistical contrasts. **Table S1.** provides an overview of the sampling techniques performed across both mainstream and sidestream WWTPs. The gaseous N_2_O emission concentrations from the BNR processes were quantified using an USEPA endorsed protocol^41^. The Surface Emission Isolated Flux Chamber (SEIFC), the only floating body approved by the Environmental Protection Agency (EPA), has a circular cross-sectional area of 0.13 m^2^ and is used to measure gaseous nitrogen fluxes from activated sludge tanks^41,42^. The SEIFC was employed to obtain headspace N_2_O concentration(ppm) and fluxes at distinct locations (aerated and nonaerated) in the treatment process sampled. At each location, online measurements of gaseous N_2_O concentration and gas flow were conducted over a 24-hour period to address the diurnal variability of gas emissions. The gaseous N_2_O concentration was measured via infrared gas filter correlation (Teledyne API, San Diego, California) at a frequency of 1/min. For emission measurements from the closed-surface, sidestream SHARON process, the N_2_O analyzer was directly connected to the off-gas stacks. Mainstream N_2_O measurements were conducted at both aerobic and anoxic zones in the target WWTPs. Measurement campaigns in both mainstream and sidestream processes were conducted over multiple days to determine the spatial and temporal variability. The use of GC-TCD in past studies to measure the off-gas flow has provided valuable data at different WWTPs around US^4,42^. We used a hotwire anemometer to directly measure the off-gas flow rates from the aerated and non-aerated zones, rather than relying on process blower data for approximating emission rates from aerated zones alone (**Supplementary section 2, Figures. S1-S3).**

**Table 1.**
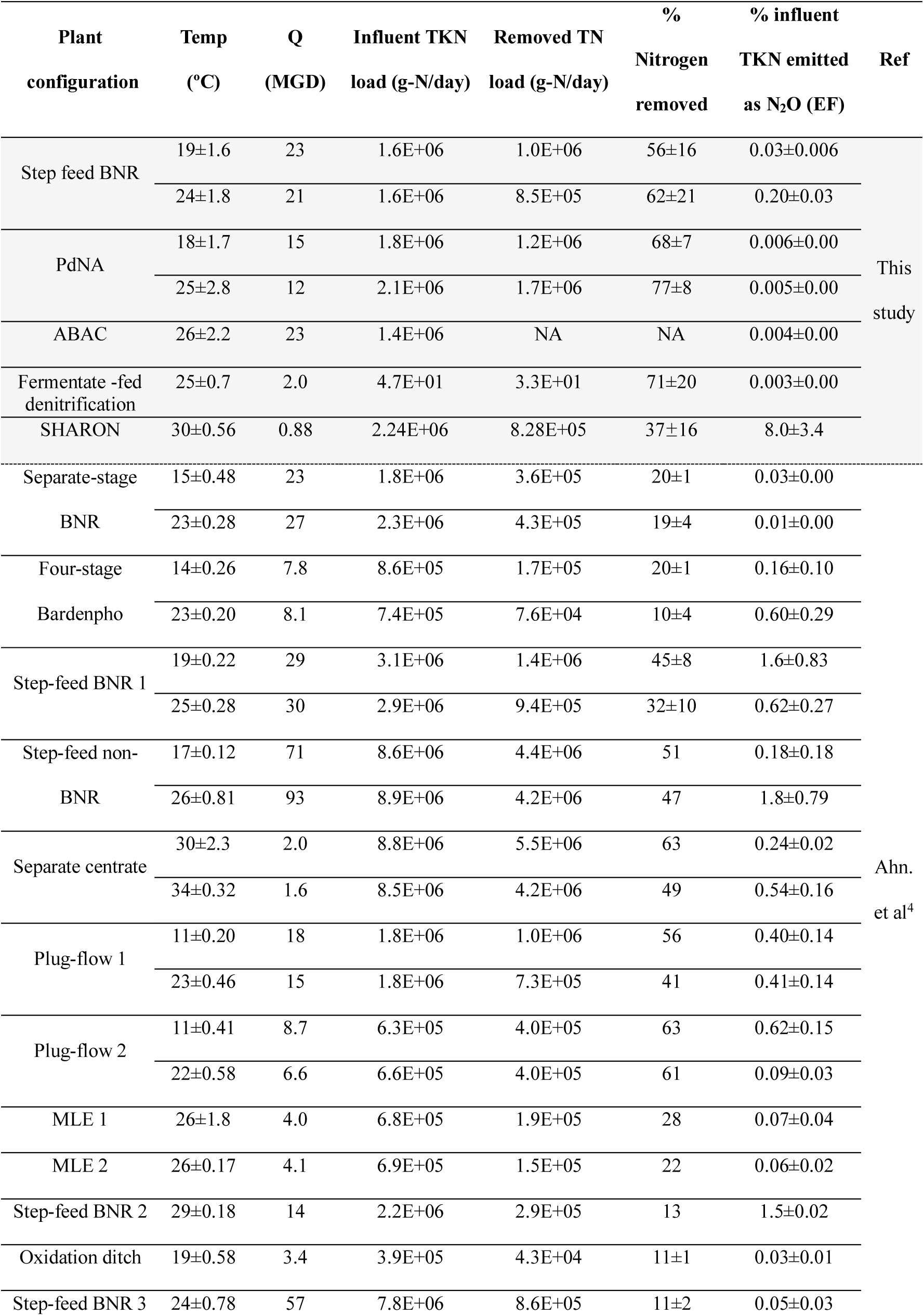
Summary of N_2_O EF from full-scale BNR WWTPs. Shaded rows indicate measurements from current study, and unshaded row represent historical data.

Concomitantly, during each monitoring campaign, wastewater nitrogen species concentrations, including influent and effluent bioreactor samples were obtained using an autosampler every 1 hour to measure NH_3_, NO_2_^-^, and NO_3_^-^ concentrations using a Lachat Quickchem Flow injection analysis system^43^. Activated sludge mixed-liquor temperature, pH, DO are acquired from the SCADA system and soluble chemical oxygen (sCOD) and total chemical oxygen demand (tCOD) were measured with Hach TNTplus kits and a Hach DR2800 spectrophotometer (HACH Loveland, CO). Mixed-liquor suspended solids (MLSS) and volatile suspended solids (VSS) concentrations were analyzed using Standard Methods 2504D and 2540E ^43^.

Bioreactor profile samples were collected from nitrifying and denitrifying tanks once a day to measure NH_3_, NO_2_^-^, and NO_3_^-^ concentrations using Hach TNTplus kits and a Hach DR2800 spectrophotometer (HACH Loveland, CO). DO concentrations, temperature (°C), and pH were measured using HACH handheld sensors (HACH Loveland, CO). Process air flow rate (standard cubic feet per minute, SCFM), SRT, the surface area of the tanks (m^2^), and influent flow rates (million gallons per day, MGD) were obtained from the plant operation records.

### 2.2. Data Preprocessing for Calculating EF and Implementing Unsupervised PCA Analysis

The measured data were first grouped into zone-specific aerobic and anoxic zone measurements to calculate N_2_O emission rates and EF across the system. The N_2_O fluxes were calculated using Eq.1, and the N_2_O emissions were calculated using Eq.2.

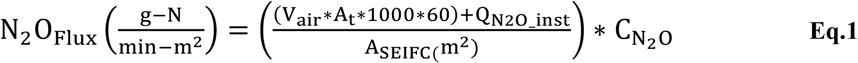

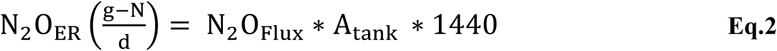

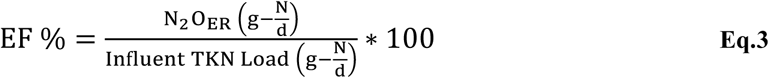

Where N_2_O_flux_ is the measure of N_2_O flux(g-N/min.m^2^) from the activated sludge tank, V_air_ is the air velocity (m.s^-1^) measured in the anemometer from the SEIFC parallel to the N_2_O concentration measurements, A_t_ is the Area of the tube(m^2^), Q_N2O_inst_ is the off-gas flow recorded on the N_2_O analyzer (L.min^-1^) from the SEIFC, A_SEIFC_ is the area of the flux chamber (m^2^) and C_N2O_ is the average concentration of the gaseous N_2_O concentration (g-N.L^-1^). N_2_O_ER_ is the N_2_O emission rate (g-N/d) from the overall tank, and A_tank_ is the area of the activated sludge tank (m^2^). To determine the Emission Factor (EF), the Q_N2O_ (N_2_O emission rate (g-N/d)) was normalized to the influent Total Kjeldahl Nitrogen (TKN) (g-N/d) load using Eq.3. All datasets used in this study were expressed to two significant figures.

For exploratory dimensionality reduction, unsupervised PCA analysis was performed separately to aerobic and anoxic datasets. The datasets consisted of both present and past datasets, which spanned different numerical ranges and units. PCA was performed using the *prcomp* function in R with variable scaling enabled (scale.=TRUE), thereby standardizing each variable to zero mean and unit variance before decomposition. This ensured that variables with larger numerical magnitudes did not disproportionately influence principal component loadings. PCA visualizations were generated using the *factoextra* package, while the Pearson correlation analysis and correlation heatmaps were developed using *corrplot* and *ggcorrplot* packages.

### 2.3. Supervised Machine Learning Approach to Predict N_2_O Fluxes

Separate from PCA, the combined past and present datasets were screened for missing and non-numeric values prior to ML analysis. To prevent data leakage, datasets were first partitioned into training (75%) and testing (25%) subsets using a stratified train-test split based on the response variables. Missing values in the training subset were imputed using the *missForest* package in R, which robustly captures complex non-linear relationships among variables^44^. Missing values in the testing subset were subsequently imputed using median statistics derived exclusively from training subset, without refitting the imputation model on test data. This workflow ensured strict separation between training and testing datasets throughout model development and evaluation.

Prior to model training, the response variables N_2_O flux, N_2_O emission rate, and N_2_O concentration were log-transformed using the *log1p* function to reduce skewness while preserving zero-valued observations. Each log-transformed response variable was subsequently min-max normalized to the [0,1] range using the scaling parameter solely from the training subset, with identical scaling applied to the testing subset. Predictor variables exhibiting substantial skewness (absolute skewness > 1) were log-transformed and substituted into the modeling dataset. The final predictor matrix consisted of influent, effluent, and operational variables from both aerobic and anoxic zones.

### 2.4. Machine Learning Models and Hyperparameter Optimization

Three ensemble tree-based supervised ML models were implemented namely Random Forest (RF), Gradient Boosting (GB), and Extreme Gradient Boosting (XGBoost). Model development and hyperparameter tuning was performed in R using the *caret*, *randomForest*, *gbm,* and *xgboost* packages.

Hyperparameter optimization was performed using 10-fold cross-validation (CV) on the training subset. For XGBoost, grid search strategy optimization was conducted over the number of boosting iterations, tree depth, learning rate, minimum child weight, and subsample ratio (**Supplementary Section 3**). Hyperparameter searches were performed for RF and GBM to identify optimal model complexity while minimizing overfitting.

### 2.5. Model Evaluation and Comparison

Model performance was evaluated using both CV and an independent held-out test dataset. CV was used to identify the optimal hyperparameter combinations for each ML model, whereas the independent held-out test dataset was used to assess the final generalization performance of the optimized models. Predictive performance was quantified using the coefficient of determination (R^2^), mean squared error (MSE), root mean square error (RMSE), and mean absolute error (MAE). To enable direct comparison with measured values, predictions generated in the normalized log-transformed space were back-transformed to their original units using reverse min-max scaling followed by the *expm1* function. All reported test-set performance metrics were calculated on the original response variable scale.

To compare the predictive performance of RF, GBM and XGBoost, repeated 10-fold CV was performed using identical resampling folds for all models. Fold-wise performance metrics were statistically compared using paired Student’s t-tests to determine whether differences in predictive accuracy between models were statistically significant.

### 2.6. Model Interpretation and Uncertainty Analysis

Model interpretation was performed to identify the operational variables governing N_2_O emissions and to evaluate whether the learned relationships were consistent with established BNR process mechanisms. Model-agnostic permutation importance was first applied to all trained models to quantify the relative contribution of each predictor by measuring the increase in prediction error following random permutation of individual variables. This provided a consistent basis for comparing variables importance across RF, GBM and XGBoost models.

To further interpret the optimized XGBoost model, Shapley Additive exPlanations (SHAP) were computed using the *iml* package with a custom prediction function. Both local and global feature contributions were evaluated. SHAP Beeswarm plots were generated using the *ggbeeswarm* and *shapviz* packages, while the *viridis* color palette was used to provide perceptually uniform color palette. Two-dimensional partial dependence plots (PDPs) were generated using the *pdp* package to characterize the marginal effects of paired operational variables on predicted N_2_O fluxes. Predictor variables were displayed in their original measurement units, while predicted responses were back-transformed from the normalized log-transformed space to the original N_2_O flux scale to facilitate physical interpretation.

To quantify model uncertainty, bootstrap resampling was performed by repeatedly retraining the optimized XGBoost model. Mechanistic consistency assessment was subsequently performed using the optimized XGBoost model to assess whether the learned response surfaces were consistent with established BNR process chemistry. Based on the dominant variables identified from the SHAP and PDP analyses, two pairwise interactions were evaluated for each operational zone: SRT-NH_3_ and DO-NH_3_, in the aerobic zone, and DO-NO_2_^-^ and SRT-DO in the anoxic zone. For each interaction, the paired variables were systematically varied across their observed ranges while all remaining predictors were fixed at their training set-median values. Model predictions were back transformed to the original N_2_O flux scale, and hotspot emission ratios were calculated by comparing predefined high and low emission operating scenarios. Bootstrap derived 95% confidence interval (CI) were then calculated for the predicted hotspot flux ratios to quantify the uncertainty associated with these mechanistic interpretations.

## 3. Results

### 3.1. Updated N_2_O emissions from advanced mainstream and sidestream activated sludge processes

We first present the results obtained from recently conducted (2023-2024) N_2_O measurement campaigns at five different next-generation mainstream and sidestream wastewater treatment processes in the Northeastern and Mid-Atlantic regions of the United States (**Table 1**). For the *<u>mainstream processes</u>* tested, the average EF, representing the fraction of the influent nitrogen load emitted as N_2_O, was lower than the IPCC EF (1.6±1.2%)^3^. Within the mainstream processes evaluated, the EF for the four-pass step-feed BNR process tested (0.20±0.03%) was higher than for the processes based on ammonia-based aeration control (ABAC) (0.004±0.00%), partial denitratation-anammox (PdNA) (0.006±0.00%), and fermentate-fed denitrification (0.003±0.00%). Based on these results by themselves, the mainstream processes sampled herein displayed lower EFs and improved N-removal relative to those previously characterized (**Table 1, Figure S4**). In contrast, the *<u>sidestream process,</u>* single reactor system for high activity ammonium removal over nitrite (SHARON), based on 47 independent daily measurements (n=47), exhibited an average EF in the range 8.0±3.4 %, which were higher than both the mainstream and IPCC-EFs.

However, some process-specific nuances could also be inferred based on the current datasets. For the mainstream PdNA process, operating temperature did not statistically impact either the BNR performance (% TN removal, which averaged 72±9 % across temperatures) or the N_2_O EF (**Table 1**). In contrast, the emissions from the step-feed BNR process were higher during the higher temperature monitoring campaign than during the lower temperature campaign (**Table 1**). On the other hand, ABAC-based process represents distinct operating characteristics with uncoupled organic carbon, nitrification, and post-denitrification, and displayed a low EF. A specific overall N-removal efficiency for the ABAC process could not be correlated with the measured N_2_O emissions, which were restricted to the aerated zones in the process (**Table 1**). Therefore, N_2_O emissions are reflective of overall process operations specific to the designed process configuration and are not strictly influenced by operating temperatures alone.

The SHARON process tested treats post-anaerobic digestion centrate and operates via a combination of nitritation (oxidation of influent NH_3_ to NO_2_^-^) followed by glycerol-mediated denitritation (reduction of NO_2_^-^ to N_2_). The SHARON process displayed low ammonia conversion (50.3±11.3%) and low nitrogen removal (37±16%) (**Table 1)**. These observed process performance measures corresponded with a higher N_2_O EF (8.0±3.4%), relative to the other processes tested **(Table 1)**. Based on previous results, sidestream treatment processes, including those operating in partial nitritation mode can display EFs as low as 0.5%^4,45^, suggesting considerable room for reduction in emissions with concomitant performance improvements in the system tested herein.

Such findings reinforce the previously demonstrated parallels between process performance and N_2_O emissions from WWTPs^7,45–47^. Nevertheless, a robust approach capable of unraveling the complex interplay between process configurations, operating conditions, process performance, and corresponding emissions is needed to explore the development of process optimization and decarbonization strategies and is described next.

### 3.2. Unraveling the Interdependencies of N_2_O Flux and Process Parameters Using a Multi-Decadal Wastewater Treatment Operations Dataset

To investigate the interdependencies between N_2_O fluxes and wastewater process variables, PCA was performed using the combined present and past wastewater process operational datasets spanning more than a decade of process monitoring (**Table 1**). First, the PCA identified the interdependencies between nitrification and denitrification process variables (nitrogen species, organic compounds, suspended solids), operating conditions (SRT, influent flow rate), and measured N_2_O fluxes. In the presented PCA biplots (**Figures 1A and 1B),** the length of the arrow represents the magnitude of the variable contribution. Positively correlated variables are clustered in the same direction as a reference variable (for instance, N_2_O fluxes), while the negative and uncorrelated variables are positioned opposite and perpendicular, respectively. We further classified the influential factors into two categories. Specifically, zone-specific variables including NH_3_, DO, NO_2_^-^ and SRT were classified as *<u>local variables</u>*, since they reflect spatially discrete sampling locations within the WWTP process bioreactors. Influent and effluent water quality parameters, influent flowrate, and temperature were considered as *<u>global variables</u>*.

**Figure 1.**
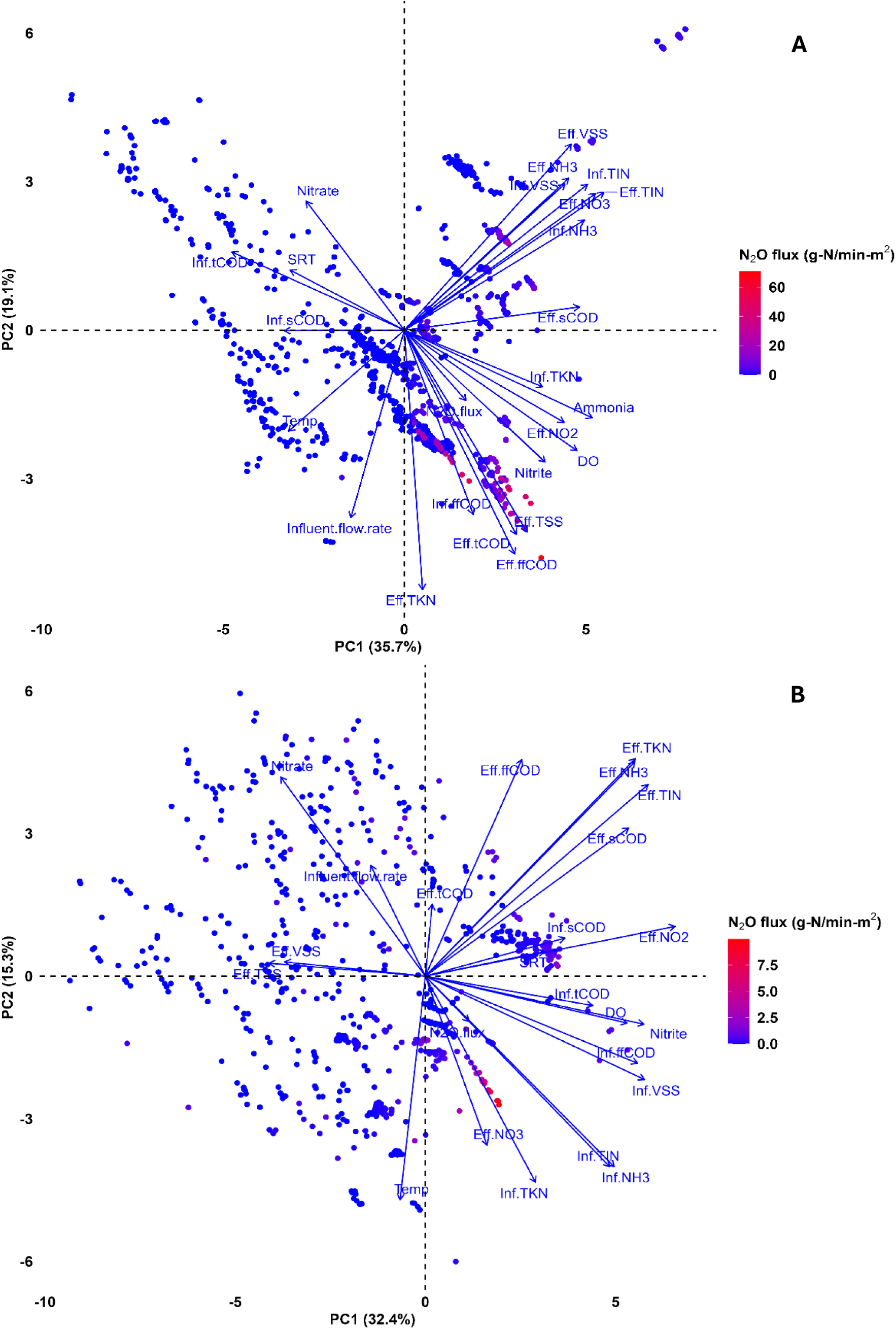
PCA analysis on N_2_O flux and wastewater quality parameters. **A, B,** Multivariate relationships between N_2_O flux (g-N min^-1^-m^2^) and local versus global variables in aerobic **(A)** and anoxic **(B)** zones. Blue arrows indicate variable loadings and their directionality along principal components, reflecting contributions to N_2_O flux.

#### 3.2.1. PCA-based interpretation of aerated zone emissions

Based on aerated zone data from current and previous sampling campaigns, PC1 and PC2 captured 35.7% and 19.1% of the variance, respectively (**Figure.1A**). Among the local variables, NH_3_, DO, NO_2_^-^ and among the global variables, influent TKN, influent ffCOD, effluent NO_2_^-^, effluent TSS, effluent ffCOD, and effluent tCOD were strongly and positively correlated with N_2_O fluxes (**Figures 1, and S5A**). One fundamental interpretation of such correlations between the local variables and observed N_2_O emissions from the aerated zones points to the role of nitrifier nitrification and nitrifier denitrification pathways. To summarize briefly, during nitrifier-nitrification, kinetic limitations in conjunction with the presence of non-limiting DO and NH_3_ concentrations in aerated zones can lead to incomplete NH_2_OH oxidation by AOB, which results in NO and N_2_O production therein^4,10,16^. Correspondingly, during nitrifier-denitrification, AOB produce N_2_O primarily under conditions that result in non-limiting NO_2_^-^ concentrations, potentially exacerbated by limiting DO and non-limiting NH_3_ concentrations^4,10^. Of the global variables, high influent TKN and effluent NO_2_^-^ concentrations could indirectly correspond with higher N_2_O fluxes depending on SRT and the corresponding extant local variables (**Figure 1A**). The observed positive correlation of measured N_2_O fluxes with different effluent TSS and organic carbon (ffCOD and tCOD) concentrations for a given degree of nitrification cannot be interpreted in isolation and needs to be considered using more complex models (such as the ML models considered below).

Negative correlations of SRT, zone-specific NO_3_^-^, and influent tCOD concentrations with measured N_2_O fluxes were also observed (**Figures 1A, and S5A**), suggesting overall that conditions that promote full nitrification (higher zone-specific NO_3_^-^ concentrations) at WWTPs (higher SRTs) correspond to lower N_2_O fluxes (in keeping with previous results)^48^. In general, the link between lower N_2_O emissions and lower influent tCOD concentrations is indirect and points to increased oxygen availability for complete nitrification compared to organic carbon oxidation^9^.

Of the remaining global variables, effluent NO_2_^-^ concentrations correlated positively with aerobic N_2_O fluxes, and likely represent the link between inadequate nitrogen removal and higher N_2_O emissions^4^ (**Figures.1A and S5A**). The global variables captured by both PC1 and PC2, including temperature, influent flow rate, influent TIN, influent NH_3_, influent VSS, effluent VSS, and effluent NH_3_, did not statistically correlate with aerobic-zone N_2_O fluxes (**Figures 1A and S5A**).

#### 3.2.2. PCA-based interpretation of anoxic-zone emissions

For data obtained from all the previously and currently sampled anoxic zones, PC1 and PC2 captured 32.4% and 15.3% of the variance, respectively. Local variables including zone-specific DO and NO_2_^-^ concentrations were positively correlated with measured N_2_O fluxes (**Figures 1B, and S5B**). Notably, zone-specific DO and NO_2_^-^ concentrations clustered closely together, emphasizing their potential significant interactive impacts on N_2_O production again linked with poor process operation and incomplete denitrification, as also shown previously ^4^.

Global variables such as effluent NO_3_^-^, effluent NO_2_^-^, influent TKN, influent NH_3_, and influent TIN concentrations exhibited a positive correlation with N_2_O fluxes (**Figures 1B and S5B**). In contrast, zone-specific NO_3_^-^ concentrations and influent flow rate were negatively correlated with N_2_O fluxes (**Figure 1B**). Effluent TKN and effluent NH_3_ concentrations were not statistically correlated with N_2_O fluxes (**Figures 1B and S5B**).

### 3.3. Hyperparameter Optimization of ML Models for Predicting N_2_O Flux under Different Wastewater Treatment Parameters

We further developed ML models with combinations of different process variables and parameters from our past and present data to predict and mitigate N_2_O emissions from WWTP operations. The ML models were trained to predict N_2_O flux (g-N/min-m^2^), N_2_O emission rate (g-N/d), and N_2_O concentration (ppm) using three ML algorithms namely RF, GB, and XGBoost. Each model was optimized under different hyperparameter tuning to identify the optimum parameter for achieving the best prediction performance (**Table S2**). Among these, XGBoost achieved the highest predictive performance of CV R^2^ of 0.98 and a held-out test R^2^ of 0.97 for aerobic N_2_O flux prediction and a CV R^2^ of 0.78 and a held-out test R^2^ of 0.87 for anoxic N_2_O flux prediction (**Table S3**).

To determine whether differences in predictive performance among models were statistically significant, repeated 10-fold CV with identical resampling folds was performed, and fold-wise performance metrics were compared using paired students t-tests. For aerobic N_2_O flux prediction, XGBoost significantly outperformed RF across R^2^, RMSE and MAE (p<0.001), whereas no statistically significant difference in R^2^ or RMSE was observed between XGBoost and GB, although XGBoost exhibited a modest but significant improvement in MAE (p<0.001). For anoxic N_2_O flux prediction, XGBoost significantly outperformed RF across R^2^ RMSE, and MAE (p<0.001), whereas no statistically significant differences in R^2^, RMSE, or MAE were observed between XGBoost and GBM (p>0.05) (**Table S3**). Therefore, XGBoost was selected for subsequent model interpretation using permutation importance, SHAP and partial dependence analyses.

### 3.4. Machine Learning Insights into the Direct Impact of Wastewater Effluent Quality on N_2_O Flux

From the trained ML models, permutation importance was extracted for both aerobic and anoxic zone conditions (**Figure 2A-C**). Of the 24 variables trained (**Figure S6**), only the top fifteen are shown in **Figure 2B, C.** The most important variables contributing to aerated-zone N_2_O emission fluxes included influent flow rate, local NH_3_, DO, and NO_2_^-^ concentrations, SRT, effluent NO_2_^-^, effluent NO_3_^-^concentrations, effluent tCOD, effluent ffCOD, and effluent TSS (**Figure 2B**). For non-aerated zones, the most important variables contributing to N_2_O emission fluxes were zone-specific DO, NO_2_^-^ and NO_3_^-^ concentrations, SRT, effluent NO_3_^-^, effluent tCOD, effluent ffCOD, effluent VSS, and effluent TIN concentrations **(Figure 2C)**. These variables identified by the ML models as influential predictors of N_2_O fluxes were consistent with the PCA analysis and underscored the now well-established link between process performance and emissions^4^. Our results also aligned with findings from a similar ML study using AdaBoost, which pointed to temperature, DO, NH_3_, NO_2_^-^, and NO_3_^-^ as being influential factors for aerobic zone N_2_O emissions, without corresponding insights into anoxic-zone emissions^27^. Our study additionally highlights the critical need to consider quantifying and predicting N_2_O emissions from anoxic zones, as their contribution to N_2_O production and emissions are inevitably under poor operating conditions^4,5^.

**Figure 2.**
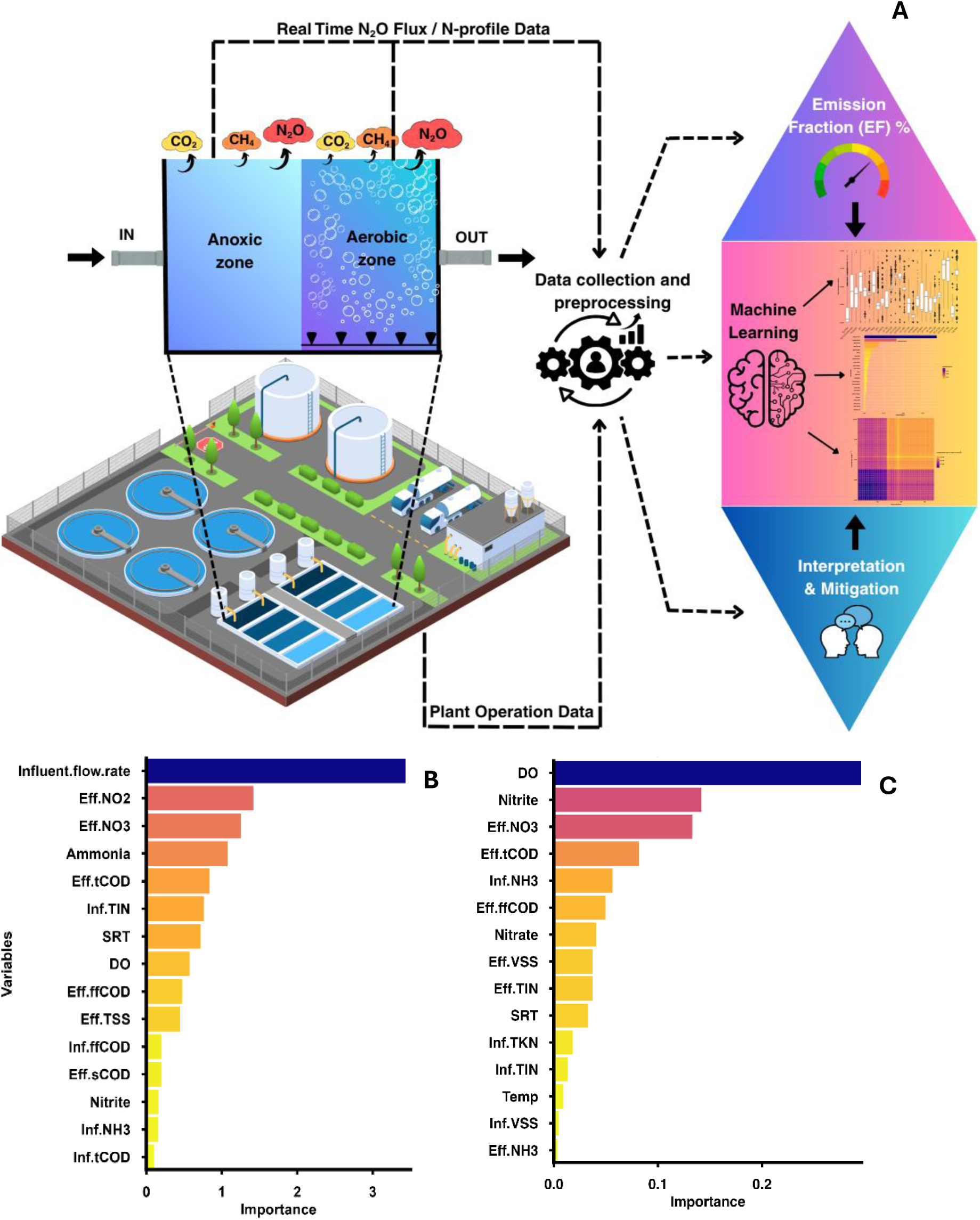
Data collection and processing in ML with variable importance from the XGBoost algorithm. **A,** workflow integrating real-time N_2_O emission data, N-profile, and plant operational data, into ML models to identify mitigation strategies to reduce N_2_O emissions and improve the water-sanitation system. **B, C,** XGBoost permutation importance scores showing contribution of individual variables to predicted N_2_O flux in aerobic **(B)** and anoxic **(C)** zones.

We further integrated SHAP analysis to uncover the magnitude and directionality influence of the variables towards predicted N_2_O flux, bridging the gap between model predictions and actionable process insights*. SHAP analysis for aerobic* N_2_O flux ranked according to mean absolute SHAP values, identified NH_3_ as the most influential predictor of aerobic N_2_O flux, followed by influent flow rate, effluent NO_3_^-^, effluent NO_2_^-^, and local NO_2_^-^ conditions (**Figure 3A**). An increase in influent flow rate and NH_3_ positively influenced predicted N_2_O flux, as reflected by their positive side of the SHAP values. Although NH_3_ exhibited the greatest overall influence, DO also demonstrated a key role. Higher NH_3_ concentrations were consistently associated with increased N_2_O flux, whereas higher DO levels could result in both higher and lower N_2_O flux, depending on the operating conditions. These results suggest that operation under non-limiting NH_3_ concentrations can promote N_2_O production, regardless of whether DO levels are high or low. Therefore, optimizing DO levels *in conjunction with* appropriate NH_3_ concentrations is essential for effective nitrification.

**Figure 3.**
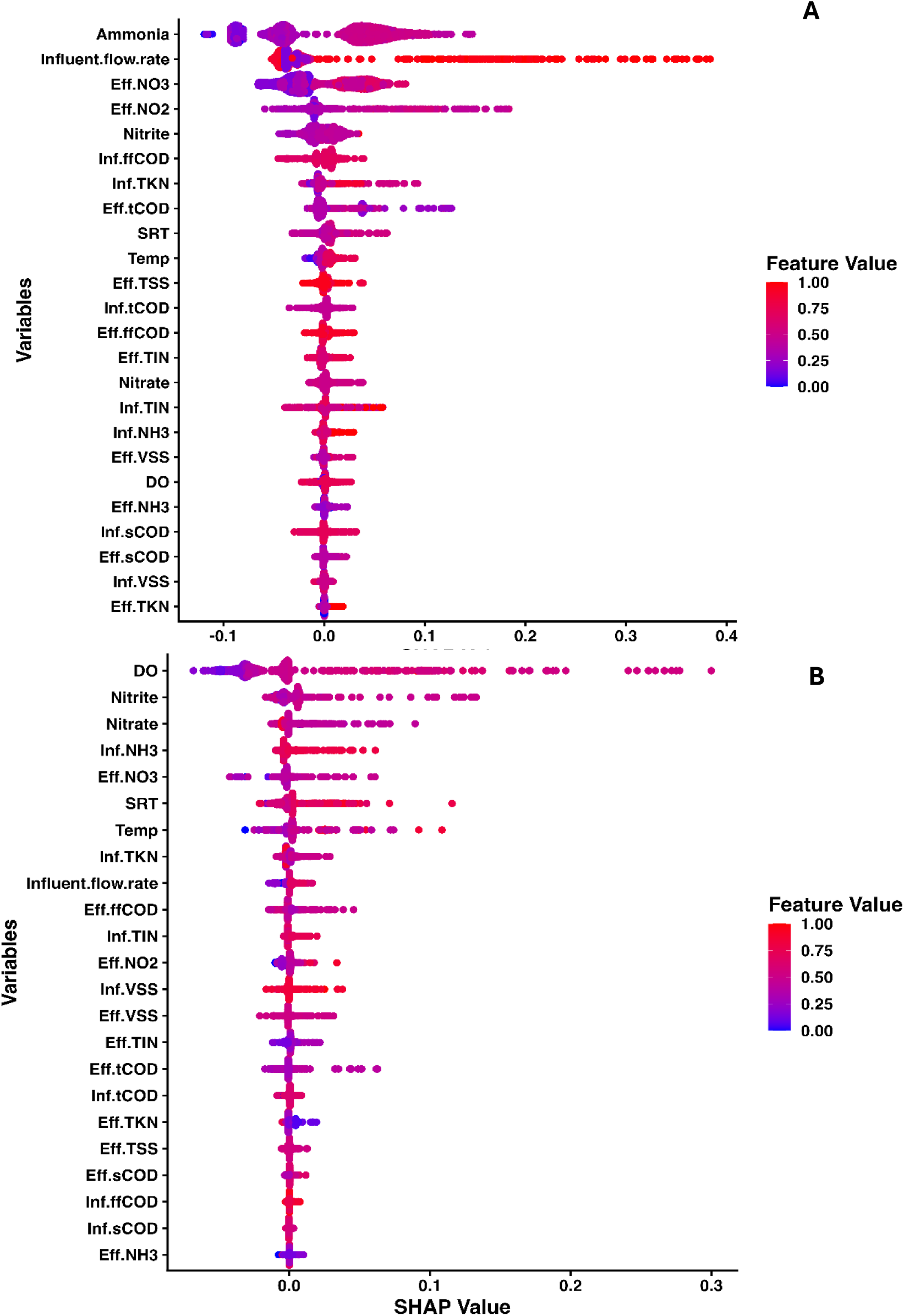
SHAP analysis of predicted N_2_O flux across wastewater parameters. **A, B,** SHAP Beeswarm plots showing the magnitude and direction of variable contributions to predicted N_2_O flux in aerobic **(A)** and anoxic **(B)** zones. The feature values are normalized to highlight variability and the influence of local and global parameters on N_2_O flux.

Our results also suggest potential interactive effects of operating SRT with DO and NH_3_ concentrations (**Figure 3A**), which have a firm biochemical basis. For instance, operation at elevated SRT can lead to reduced N_2_O fluxes, coupled with optimization of DO concentrations to achieve a given NH_3_ conversion. Based on the mean absolute SHAP values, NO_2_^-^ concentrations exhibited a greater overall influence on N_2_O fluxes compared to that of DO and SRT. However, the production of NO_2_^-^ can only be controlled by concurrently optimizing DO and SRT along with NH_3_ concentration.

#### SHAP analysis for anoxic

N_2_O flux ranked DO as the most influential predictor based on mean absolute SHAP values, followed by local NO_2_^-^, local NO_3_^-^, influent NH_3_, and effluent NO_3_^-^ concentrations (**Figure 3B**). Higher DO on the positive side could be linked to DO-mediated inhibition of N_2_O reduction to N ^4,5,18^. A biphasic response of N_2_O fluxes to SRT values was observed once again pointing to possible interactions with extant DO and NO_2_^-^ concentrations.

Although NO_3_^-^ ranked among the most influential predictors, its effect was smaller than that DO and NO_2_^-^. Higher NO_3_^-^ feature value representing positive SHAP values (**Figure 3B**) does not directly imply its involvement in producing N_2_O. For both aerobic and anoxic zones, the global variables, effluent NO_2_^-^ and NO_3_^-^ have higher feature values in the positive side of the SHAP values (**Figure 3A, B**), reflecting that higher effluent NO_2_^-^ and NO_3_^-^ concentrations are associated with increased predicted N_2_O flux, as it represents an incomplete nitrification and ineffective denitrification process. In addition, SHAP analysis for N_2_O emission rate and N_2_O concentration under both aerobic and anoxic zones are provided in **Figure S7,** further supporting these trends.

### 3.5. PDP-derived insights on operational regions of low predicted N_2_O flux

#### 3.5.1. Aerobic zone

Two-dimensional PDP analysis was applied to characterize the predicted response of N_2_O flux across paired operational variables, with SRT considered alongside NH_3_ and DO as the principal process parameters of interest. **Figure 4A** indicates that NH_3_ concentrations below 3 mg L^-1^ at SRT below 10 days correspond to lower predicted N_2_O flux. The DO and NH_3_ interaction (**Figure 4B**) show that operating regions characterized by NH_3_ concentrations below 3 mg L^-1^ together with DO concentrations below 4 mg O_2_ L^-1^ are associated with lower model predicted N_2_O flux. In contrast, operating conditions that foster concomitant non-limiting DO and NH_3_ conditions correspond to elevated predicted N_2_O fluxes. (**Figure 4B**). Such predicted responses are in excellent alignment with fundamental and mechanistic drivers of N_2_O emissions under aerobic conditions in WWTPs, including combined contributions of oxidative and reductive pathways primarily in AOB^4–9,45–49^.

**Figure 4.**
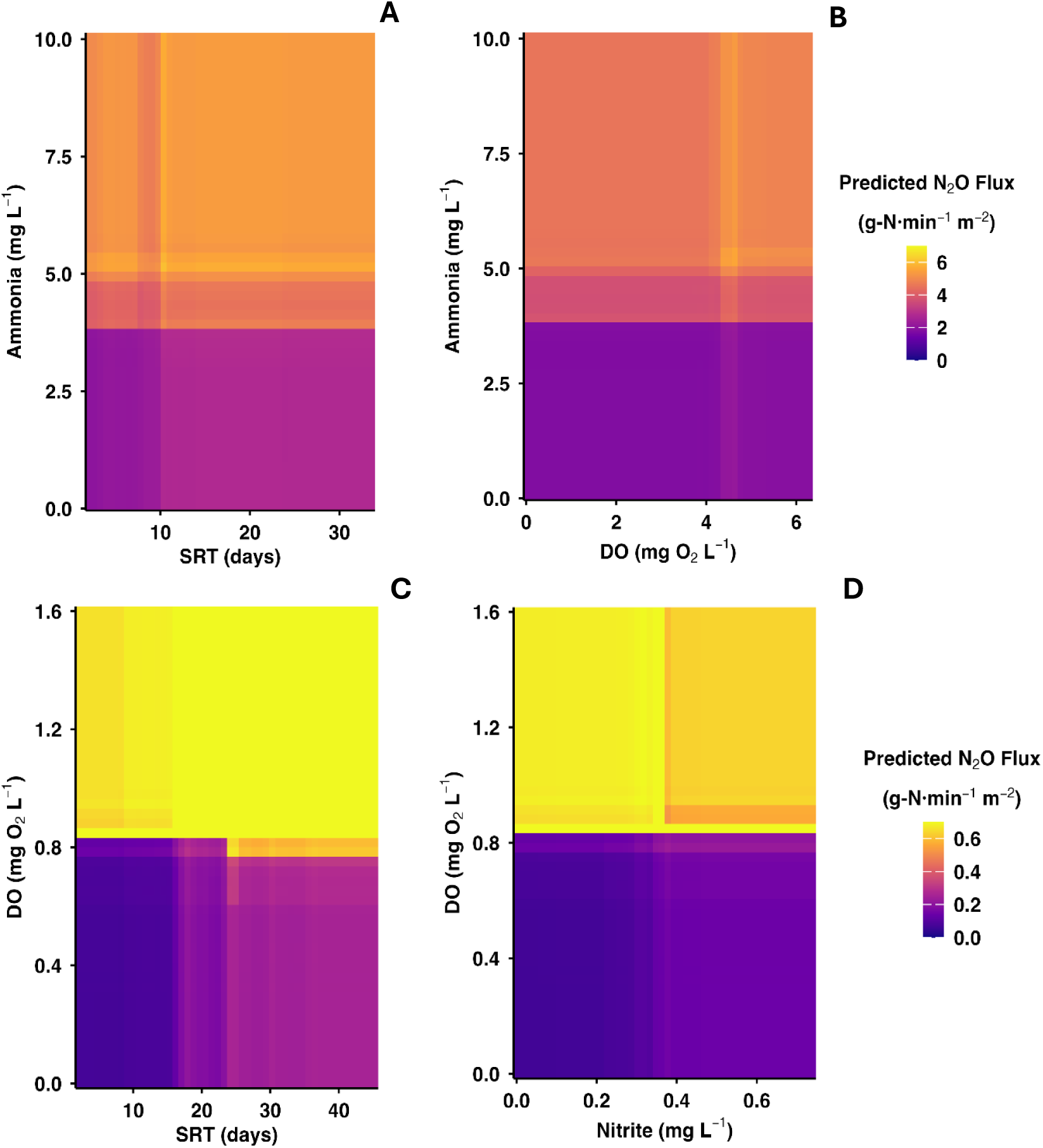
PDP on predicted N_2_O flux with selective key features. Two-dimensional PDP showing predicted N_2_O flux and its dynamic relationship towards key process variables in aerobic **(A, B)** and anoxic **(C, D)** zones. Predicted N_2_O flux with **(A)** NH_3_ and SRT, **(B)** NH_3_ and DO, **(C)** DO and SRT, **(D)** DO and NO_2_^-^.

#### 3.5.2. Anoxic zone

Similarly, conditions that show lower predicted N_2_O fluxes were associated with DO concentrations below 1 mg O_2_ L^-1^ coupled with total process SRTs below 20 days (**Figure 4C**). Elevated predicted N_2_O fluxes at DO concentrations above 1 mg O_2_ L^-1^ and SRTs below 20 days (**Figure 4C**) coincide with NO_2_^-^ accumulation (**Figure 4D**), and concurrent non-limiting DO and NO_2_^-^ concentrations correspond to elevated predicted N_2_O fluxes even at SRTs above 20 days (**Figure 4C, D**. In a more general sense, we acknowledge that operation at such high SRT values may not be an option for typical WWTPs that employ suspended-phase activated sludge processes. DO control emerges as the dominant operational variable associated with reduced predicted NO_2_^-^ accumulation and N_2_O flux in the anoxic zone, while elevated SRTs may provide a secondary lever where feasible.

### 3.6. Mechanistic Consistency Assessment of the Learned N_2_O Response Surface

The trained model was evaluated on two-dimensional scenario grids to assess whether the learned response surface was consistent with established mechanisms of biological N_2_O production (**Figure 4**) (**Table S4**). Bootstrap uncertainty was quantified using 95% CI for each predicted scenario ratio. For the *aerobic zone*, the evaluated interactions produced predicted flux ratios consistent with established AOB-mediated pathways of N_2_O production (**Figure 4A,B**) (**Table S4**). The SRT - NH_3_ interaction produced the largest predicted contrast, with the median predicted N_2_O fluxes were 8.91-fold higher under short SRT and high NH_3_ conditions (3.22 to 38.72-fold) (**Figure 4A**) (**Table S4**). This is consistent with the SRT dependent maturity of the nitrifying community. Short SRTs near AOB washout allow NH_3_ accumulation and partial oxidation, whereas longer SRTs support stable and complete nitrification^9^. The DO - NH_3_ interaction produced the second largest predicted contrast, with median predicted N_2_O fluxes were 7.83-fold higher under low DO and high NH_3_ conditions (2.22 to 52.82-fold) (**Figure 4B**) (**Table S4**). This is consistent with AOB shifting from complete NH_3_ oxidation toward partial reduction of NO_2_^-^ to N_2_O via nitrifier denitrification pathway under low DO and high NH_3_ conditions^6–8^.

For the *anoxic zone*, two tested interactions produced ratios are consistent with established denitrification biochemistry (**Figure 4C,D**) (**Table S4**). The DO-NO_2_^-^ interaction produced the largest predicted contrast, with median predicted N_2_O fluxes were 8.80-fold higher under high DO and high NO_2_^-^ conditions (3.83 to 21.33-fold) (**Figure 4D**) (**Table S4**). This is consistent with the well-documented inhibition of Nos reductase activity by residual oxygen and accumulating NO_2_^-^, which suppresses the final reductive step of denitrification and promotes N_2_O production^5,9,11^. The DO - SRT interaction produced the second largest predicted contrast, with median predicted N_2_O fluxes were 5.15-fold higher under high DO and short SRT conditions (1.47 to 18.80-fold) **(Figure 4C) (Table S4**), reflecting the secondary modulating role of SRT in denitrification performance under oxygen stress.

## 4. Discussion

### 4.1. Poor process operation leads to high N_2_O emissions irrespective of process configuration and temperature variations

Based on a combination of current measurements and extensive historical datasets, we determined that less than optimal process operations can manifest themselves in combinations of both local and global variables that in turn are correlated with higher N_2_O production and emission across aerobic and anoxic zones. Among the identified variables, SRT and DO emerge as the critical control factors associated with emissions. The key local and global variables identified from the unsupervised PCA followed by supervised ML permutation importance and SHAP aligned well with the established understanding of the nitrification and denitrification processes^4–7,29–31^, thus providing consistent insights into the contribution and significance linking to the process operation, N_2_O emission, and effluent water quality. The PDP analysis conducted for aerobic zones illustrates the relationship of NH_3_ concentration with key process control variables, DO and SRT, identifying the operating regions associated with effective nitrification and reduced predicted N_2_O flux (**Figure 4A, B**). Similarly, in anoxic zones, the PDP analysis highlights the interactions among DO, SRT, and NO_2_^-^ pinpointing operational regions associated with achieving effective denitrification and reduced predicted N_2_O flux (**Figure 4C, D**). The mechanistic consistency assessment extended this interpretation by quantitatively evaluating whether the model’s directional responses were consistent with established BNR biochemistry. Four of the tested variable interactions produced predicted flux ratios consistent with nitrifier denitrification in the aerobic zone and the DO - NO_2_^-^ mediated inhibition of N_2_O reductase activity in the anoxic zone. Thus, these ML-based interpretations reveal the interplay between the operational variables and biochemical fundamentals, that contribute to N_2_O fluxes. Nevertheless, such analysis provides a defensible basis for identifying operating regions for reducing both *aqueous* and *gaseous* nitrogenous discharge.

### 4.2. Integration of real-time process control based on ML framework

This study illustrates the potential of data-driven models, interpreted in conjunction with bioprocess fundamentals, for supporting process control and optimization in BNR systems. Upon further integrating such models into Supervisory Control And Data Acquisition (SCADA) systems, they continuously process data generated from real-time or non-real-time measurements, facilitating the development of digital twin architectures based on model-based systems engineering (MBSE)^50^. Data-driven models can provide early fault detection and advanced process control capabilities, which mechanistic models typically cannot deliver using real-time data^50^. Indeed, previous studies have successfully applied data-driven approaches in real-time reactor operations, enhancing reactor performance and effluent water quality^51–54^. The emphasis on data-driven modeling for GHG emissions, particularly N_2_O emissions, continues to expand, driven by ongoing efforts to better understand process dynamics better and improve forecasting accuracy^27,37–40^. As such, substantial opportunities exist to integrate advanced interpretation tools, including PDP and mechanistic consistency assessment, as demonstrated herein, with digital and hybrid twin systems for optimizing WWTP operations, including managing their GHG footprint.

### 4.3. Limitations

Despite the strong predictive performance achieved in this study, some limitations should be acknowledged. Although the model was developed using much time-resolved observations, the range of independent operating conditions represented in the training data remained limited, which may affect its generalizability to other WRRFs. Consequently, while model performance was evaluated using an internal hold-out dataset and repeated CV, validation using independent external datasets remains an important next step. Furthermore, the interpretability analyses describe relationships learned by the XGBoost model and should therefore be viewed as mechanistic consistency with current process understanding rather than direct evidence of causal biological mechanisms. Uncertainty associated with flux chamber N_2_O flux measurements was also not explicitly propagated through the ML framework, and the omission of process configuration as a predictor may have partially confounded the learned temperature effects.

Future work will focus on expanding the dataset across multiple facilities and operating conditions, incorporating external validation and uncertainty propagation, and integrating additional operational parameters and microbial characterization to further improve model robustness, interpretability, and generalization.

## 5. Conclusions

N_2_O emissions from BNR processes remain a significant contributor to the carbon footprint of wastewater treatment and identifying the operational strategies that concurrently minimize gaseous and aqueous nitrogenous discharge is an ongoing priority. This study integrated 2023-2024 N_2_O measurement campaigns at five next generation BNR processes with a layered ML interpretation framework applied to a multi-decadal operational dataset. Across diverse process configurations and operating temperatures, next generation mainstream BNR configurations including ABAC and PdNA achieved EF more than two orders of magnitude lower than the IPCC EF, while step-feed and sidestream processes operated under suboptimal conditions exhibited substantially higher emissions. The supervised ML interpretation identified DO, SRT, NH_3_ and NO_2_^-^ as the dominant operational variables associated with predicted N_2_O flux and identified operating regions of lower predicted flux centered at DO < 4 mg O_2_ L^-1^ in aerobic zones and < 1 mg O_2_ L^-1^ in anoxic zones, coupled with SRTs supporting stable nitrification and denitrification. Mechanistic consistency assessment further demonstrated that the learned response surfaces were consistent with established biological pathways governing N_2_O production. In summary, our results demonstrated that interpretable ML grounded in biochemical fundamentals can identify operating regions associated with reduced N_2_O emissions and provide a foundation for data-driven optimization of BNR systems.

## References

(1) Ravishankara, A. R.; Daniel, J. S.; Portmann, R. W. Nitrous Oxide (N_2_O): The Dominant Ozone-Depleting Substance Emitted in the 21st Century. Science 2009, 326 (5949), 123–125.

(2) Stocker, T. Climate Change 2013: The Physical Science Basis: Working Group I Contribution to the Fifth Assessment Report of the Intergovernmental Panel on Climate Change; Cambridge University Press, 2014.

(3) 2019 Refinement to the 2006 IPCC Guidelines for National Greenhouse Gas Inventories — IPCC. https://www.ipcc.ch/report/2019-refinement-to-the-2006-ipcc-guidelines-for-national-greenhouse-gas-inventories/ (accessed 2024-02-26).

(4) Ahn, J. H.; Kim, S.; Park, H.; Rahm, B.; Pagilla, K.; Chandran, K. N2O Emissions from Activated Sludge Processes, 2008-2009: Results of a National Monitoring Survey in the United States. Environmental Science and Technology 2010, 44 (12), 4505–4511. 10.1021/es903845y.

(5) Rassamee, V.; Sattayatewa, C.; Pagilla, K.; Chandran, K. Effect of Oxic and Anoxic Conditions on Nitrous Oxide Emissions from Nitrification and Denitrification Processes. Biotechnology and Bioengineering 2011, 108 (9), 2036–2045. 10.1002/bit.23147.

(6) Yu, R.; Kampschreur, M. J.; Loosdrecht, M. C. M. van; Chandran, K. Mechanisms and Specific Directionality of Autotrophic Nitrous Oxide and Nitric Oxide Generation during Transient Anoxia. Environ. Sci. Technol. 2010, 44 (4), 1313–1319. 10.1021/es902794a.

(7) Brotto, A. C.; Li, H.; Dumit, M.; Gabarró, J.; Colprim, J.; Murthy, S.; Chandran, K. Characterization and Mitigation of Nitrous Oxide (N2O) Emissions from Partial and Full-Nitrification BNR Processes Based on Post-Anoxic Aeration Control. Biotechnology and Bioengineering 2015, 112 (11), 2241–2247. 10.1002/bit.25635.

(8) Kampschreur, M. J.; Temmink, H.; Kleerebezem, R.; Jetten, M. S. M.; van Loosdrecht, M. C. M. Nitrous Oxide Emission during Wastewater Treatment. Water Research 2009, 43 (17), 4093–4103. 10.1016/j.watres.2009.03.001.

(9) Grady, C. P. L. Biological Wastewater Treatment; New York: Marcel Dekker, 1999.

(10) Chandran, K.; Stein, L. Y.; Klotz, M. G.; Van Loosdrecht, M. C. M. Nitrous Oxide Production by Lithotrophic Ammonia-Oxidizing Bacteria and Implications for Engineered Nitrogen-Removal Systems. Biochemical Society Transactions 2011, 39 (6), 1832–1837. 10.1042/BST20110717.

(11) Zumft, W. G. Cell Biology and Molecular Basis of Denitrification. Microbiol Mol Biol Rev 1997, 61 (4), 533–616. 10.1128/mmbr.61.4.533-616.1997.

(12) Kester, R. A.; De Boer, W.; Laanbroek, H. J. Production of NO and N(Inf2)O by Pure Cultures of Nitrifying and Denitrifying Bacteria during Changes in Aeration. Appl Environ Microbiol 1997, 63 (10), 3872–3877.

(13) Ritchie, G. A.; Nicholas, D. J. Identification of the Sources of Nitrous Oxide Produced by Oxidative and Reductive Processes in Nitrosomonas Europaea. Biochem J 1972, 126 (5), 1181–1191. 10.1042/bj1261181.

(14) Stüven, R.; Vollmer, M.; Bock, E. The Impact of Organic Matter on Nitric Oxide Formation by Nitrosomonas Europaea. Arch. Microbiol. 1992, 158 (6), 439–443. 10.1007/BF00276306.

(15) Yu, R.; Perez-Garcia, O.; Lu, H.; Chandran, K. *Nitrosomonas Europaea* Adaptation to Anoxic-Oxic Cycling: Insights from Transcription Analysis, Proteomics and Metabolic Network Modeling. Science of The Total Environment 2018, 615, 1566–1573. 10.1016/j.scitotenv.2017.09.142.

(16) Schreiber, F.; Wunderlin, P.; Udert, K. M.; Wells, G. F. Nitric Oxide and Nitrous Oxide Turnover in Natural and Engineered Microbial Communities: Biological Pathways, Chemical Reactions, and Novel Technologies. Front Microbiol 2012, 3, 372. 10.3389/fmicb.2012.00372.

(17) Wunderlin, P.; Mohn, J.; Joss, A.; Emmenegger, L.; Siegrist, H. Mechanisms of N2O Production in Biological Wastewater Treatment under Nitrifying and Denitrifying Conditions. Water Research 2012, 46 (4), 1027–1037. 10.1016/j.watres.2011.11.080.

(18) Lu, H.; Chandran, K. Factors Promoting Emissions of Nitrous Oxide and Nitric Oxide from Denitrifying Sequencing Batch Reactors Operated with Methanol and Ethanol as Electron Donors. Biotechnology and Bioengineering 2010, 106 (3), 390–398. 10.1002/bit.22704.

(19) US EPA, O. Effluent Guidelines. https://www.epa.gov/eg (accessed 2025-07-28).

(20) Daelman, M. R. J.; van Voorthuizen, E. M.; van Dongen, U. G. J. M.; Volcke, E. I. P.; van Loosdrecht, M. C. M. Seasonal and Diurnal Variability of N2O Emissions from a Full-Scale Municipal Wastewater Treatment Plant. Science of the Total Environment 2015, 536, 1–11. 10.1016/j.scitotenv.2015.06.122.

(21) Gernaey, K. V.; van Loosdrecht, M. C. M.; Henze, M.; Lind, M.; Jørgensen, S. B. Activated Sludge Wastewater Treatment Plant Modelling and Simulation: State of the Art. Environmental Modelling & Software 2004, 19 (9), 763–783. 10.1016/j.envsoft.2003.03.005.

(22) Perez-Garcia, O.; Mankelow, C.; Chandran, K.; Villas-Boas, S. G.; Singhal, N. Modulation of Nitrous Oxide (N2O) Accumulation by Primary Metabolites in Denitrifying Cultures Adapting to Changes in Environmental C and N. Environ. Sci. Technol. 2017, 51 (23), 13678–13688. 10.1021/acs.est.7b03345.

(23) Frontiers | Metabolic Network Modeling of Microbial Interactions in Natural and Engineered Environmental Systems. https://www.frontiersin.org/journals/microbiology/articles/10.3389/fmicb.2016.00673/full (accessed 2025-03-17).

(24) Perez-Garcia, O.; Chandran, K.; Villas-Boas, S. G.; Singhal, N. Assessment of Nitric Oxide (NO) Redox Reactions Contribution to Nitrous Oxide (N2O) Formation during Nitrification Using a Multispecies Metabolic Network Model. Biotechnology and Bioengineering 2016, 113 (5), 1124–1136. 10.1002/bit.25880.

(25) Perez-Garcia, O.; Villas-Boas, S. G.; Swift, S.; Chandran, K.; Singhal, N. Clarifying the Regulation of NO/N2O Production in *Nitrosomonas Europaea* during Anoxic–Oxic Transition via Flux Balance Analysis of a Metabolic Network Model. Water Research 2014, 60, 267–277. 10.1016/j.watres.2014.04.049.

(26) Behera, C. R.; Srinivasan, B.; Chandran, K.; Venkatasubramanian, V. Model Based Predictive Control for Energy Efficient Biological Nitrification Process with Minimal Nitrous Oxide Production. Chemical Engineering Journal 2015, 268, 300–310. 10.1016/j.cej.2015.01.044.

(27) Khalil, M.; AlSayed, A.; Liu, Y.; Vanrolleghem, P. A. Machine Learning for Modeling N2O Emissions from Wastewater Treatment Plants: Aligning Model Performance, Complexity, and Interpretability. Water Research 2023, 245, 120667. 10.1016/j.watres.2023.120667.

(28) Bellandi, G.; Weijers, S.; Gori, R.; Nopens, I. Towards an Online Mitigation Strategy for N2O Emissions through Principal Components Analysis and Clustering Techniques. J Environ Manage 2020, 261, 110219. 10.1016/j.jenvman.2020.110219.

(29) Villez, K.; Sin, G.; Vanrolleghem, P. A.; Ruiz, M.; Colomer, J.; Rosén, C.; Vanrolleghem, P. A. Combining Multiway Principal Component Analysis (MPCA) and Clustering for Efficient Data Mining of Historical Data Sets of SBR Processes. Water Sci Technol 2008, 57 (10), 1659–1666. 10.2166/wst.2008.143.

(30) Sobczyk, M.; Pajdak-Stós, A.; Fiałkowska, E.; Sobczyk, Ł.; Fyda, J. Multivariate Analysis of Activated Sludge Community in Full-Scale Wastewater Treatment Plants. Environ Sci Pollut Res 2021, 28 (3), 3579–3589. 10.1007/s11356-020-10684-5.

(31) Cui, F.; Kim, M.; Park, C.; Kim, D.; Mo, K.; Kim, M. Application of Principal Component Analysis (PCA) to the Assessment of Parameter Correlations in the Partial-Nitrification Process Using Aerobic Granular Sludge. Journal of Environmental Management 2021, 288, 112408. 10.1016/j.jenvman.2021.112408.

(32) Çimen Mesutoğlu, Ö.; Gök, O. Prediction of COD in Industrial Wastewater Treatment Plant Using an Artificial Neural Network. Sci Rep 2024, 14 (1), 13750. 10.1038/s41598-024-64634-z.

(33) Liu, T.; Zhang, H.; Wu, J.; Liu, W.; Fang, Y. Wastewater Treatment Process Enhancement Based on Multi-Objective Optimization and Interpretable Machine Learning. Journal of Environmental Management 2024, 364, 121430. 10.1016/j.jenvman.2024.121430.

(34) Mathur, R.; Sharma, M. K.; Loganathan, K.; Abbas, M.; Hussain, S.; Kataria, G.; Alqahtani, M. S.; Srinivas Rao, K. Modeling of Two-Stage Anaerobic Onsite Wastewater Sanitation System to Predict Effluent Soluble Chemical Oxygen Demand through Machine Learning. Sci Rep 2024, 14 (1), 1835. 10.1038/s41598-023-50805-x.

(35) Zhang, Y.; Wu, H.; Xu, R.; Wang, Y.; Chen, L.; Wei, C. Machine Learning Modeling for the Prediction of Phosphorus and Nitrogen Removal Efficiency and Screening of Crucial Microorganisms in Wastewater Treatment Plants. Science of The Total Environment 2024, 907, 167730. 10.1016/j.scitotenv.2023.167730.

(36) Sheik, A. G.; Malla, M. A.; Srungavarapu, C. S.; Patan, A. K.; Kumari, S.; Bux, F. Prediction of Wastewater Quality Parameters Using Adaptive and Machine Learning Models: A South African Case Study. Journal of Water Process Engineering 2024, 67, 106185. 10.1016/j.jwpe.2024.106185.

(37) Seshan, S.; Poinapen, J.; Zandvoort, M. H.; van Lier, J. B.; Kapelan, Z. Forecasting Nitrous Oxide Emissions from a Full-Scale Wastewater Treatment Plant Using LSTM-Based Deep Learning Models. Water Research 2025, 268, 122754. 10.1016/j.watres.2024.122754.

(38) Xu, X.; Wei, A.; Tang, S.; Liu, Q.; Shi, H.; Sun, W. Prediction of Nitrous Oxide Emission of a Municipal Wastewater Treatment Plant Using LSTM-Based Deep Learning Models. Environ Sci Pollut Res 2024, 31 (2), 2167–2186. 10.1007/s11356-023-31250-9.

(39) Li, K.; Duan, H.; Liu, L.; Qiu, R.; van den Akker, B.; Ni, B. J.; Chen, T.; Yin, H.; Yuan, Z.; Ye, L. An Integrated First Principal and Deep Learning Approach for Modeling Nitrous Oxide Emissions from Wastewater Treatment Plants. Environmental Science and Technology 2022, 56 (4), 2816–2826. 10.1021/acs.est.1c05020.

(40) Khalil, M.; AlSayed, A.; Liu, Y.; Vanrolleghem, P. A. An Integrated Feature Selection and Hyperparameter Optimization Algorithm for Balanced Machine Learning Models Predicting N2O Emissions from Wastewater Treatment Plants. Journal of Water Process Engineering 2024, 63, 105512. 10.1016/j.jwpe.2024.105512.

(41) Chandran, K.; Pagilla, K.; Katehis, D. Field Protocol With Quality Assurance Plan - Characterization of Nitrogen Greenhouse Gas Emissions From Wastewater Treatment Bnr Operations. 2009, 1–47.

(42) Chandran, K. Protocol for the Measurement of Nitrous Oxide Fluxes from Biological Wastewater Treatment Plants; Elsevier Inc., 2011; Vol. 486. 10.1016/B978-0-12-381294-0.00016-X.

(43) Andrew D. Eaton, American Water Works Association, Water Environment Federation. Standard Methods for the Examination of Water and Wastewater; APHA-AWWA-WEF, Washington, D.C., 2005.

(44) Stekhoven, D. J.; Bühlmann, P. MissForest—Non-Parametric Missing Value Imputation for Mixed-Type Data. Bioinformatics 2012, 28 (1), 112–118. 10.1093/bioinformatics/btr597.

(45) Ahn, J. H.; Kwan, T.; Chandran, K. Comparison of Partial and Full Nitrification Processes Applied for Treating High-Strength Nitrogen Wastewaters: Microbial Ecology through Nitrous Oxide Production. Environ. Sci. Technol. 2011, 45 (7), 2734–2740. 10.1021/es103534g.

(46) Kim, Y. M.; Park, H.; Chandran, K. Nitrification Inhibition by Hexavalent Chromium Cr(VI)--Microbial Ecology, Gene Expression and off-Gas Emissions. Water Res 2016, 92, 254–261. 10.1016/j.watres.2016.01.042.

(47) Brotto, A. C.; Annavajhala, M. K.; Chandran, K. Metatranscriptomic Investigation of Adaptation in NO and N2O Production From a Lab-Scale Nitrification Process Upon Repeated Exposure to Anoxic–Aerobic Cycling. Front Microbiol 2018, 9, 3012. 10.3389/fmicb.2018.03012.

(48) Li, B.; Wu, G. Effects of Sludge Retention Times on Nutrient Removal and Nitrous Oxide Emission in Biological Nutrient Removal Processes. Int J Environ Res Public Health 2014, 11 (4), 3553–3569. 10.3390/ijerph110403553.

(49) Brotto, A. C.; Kligerman, D. C.; Andrade, S. A.; Ribeiro, R. P.; Oliveira, J. L. M.; Chandran, K.; de Mello, W. Z. Factors Controlling Nitrous Oxide Emissions from a Full-Scale Activated Sludge System in the Tropics. Environ Sci Pollut Res 2015, 22 (15), 11840–11849. 10.1007/s11356-015-4467-x.

(50) Wang, A.-J.; Li, H.; He, Z.; Tao, Y.; Wang, H.; Yang, M.; Savic, D.; Daigger, G. T.; Ren, N. Digital Twins for Wastewater Treatment: A Technical Review. Engineering 2024, 36, 21–35. 10.1016/j.eng.2024.04.012.

(51) Zhou, J.; Huang, F.; Shen, W.; Liu, Z.; Corriou, J.-P.; Seferlis, P. Sub-Period Division Strategies Combined with Multiway Principle Component Analysis for Fault Diagnosis on Sequence Batch Reactor of Wastewater Treatment Process in Paper Mill. Process Safety and Environmental Protection 2021, 146, 9–19. 10.1016/j.psep.2020.08.032.

(52) AL-Kordy, S. U.; Khudair, D. B. H. Effluent Quality Assessment of Sewage Treatment Plant Using Principal Component Analysis and Cluster Analysis. Journal of Engineering 2021, 27 (4), 79–95. 10.31026/j.eng.2021.04.07.

(53) Liu, H.; Yang, J.; Zhang, Y.; Yang, C. Monitoring of Wastewater Treatment Processes Using Dynamic Concurrent Kernel Partial Least Squares. Process Safety and Environmental Protection 2021, 147, 274–282. 10.1016/j.psep.2020.09.034.

(54) Yang, C.; Zhang, Y.; Huang, M.; Liu, H. Adaptive Dynamic Prediction of Effluent Quality in Wastewater Treatment Processes Using Partial Least Squares Embedded with Relevance Vector Machine. Journal of Cleaner Production 2021, 314, 128076. 10.1016/j.jclepro.2021.128076.

