## Supplemental file for "Machine learning informs mitigation strategies for nitrous oxide emissions from wastewater operations"

### Section 1: Summary of wastewater sampling at WWTPs

**Table S1. Summary of sampling procedures for gaseous nitrous oxide and process operating conditions at different WWTPs**

| Type | Details | Measurement methods | Data frequency |
|---|---|---|---|
| $N_2O$ concentration using gas-phase analyser | Real-time $N_2O$ concentration | SEIFC and Teledyne $N_2O$ analyser[2] | Every 1 min [1,2] |
| Airflow measurement | Measure the airflow coming out from SEIFC | Anemometer (Extech SDL350) Hotwire thermo anemometer and datalogger | Every 1 min |
| Influent/ Effluent wastewater characteristics | $NH_3$, $NO_2^-$, $NO_3^-$, COD, TSS, VSS | Flow injection analysis, Hach TNTplus kits and 2504D and 2540E method[3] | Twice a week, every 1 hour a day (autosampler) |
| Liquid flow rate and air flow rate | Wastewater flowrate and Air blower data | SCADA | Every 1 hour a day |
| Profile sampling at liquid phase | $NH_3$, $NO_2^-$, $NO_3^-$, DO, Temperature and pH | Hach TNTplus kits and Hach DR2800 spectrophotometer, and Hach handheld sensors (Hach Loveland, CO) | Once a day (grab sample) |

### Section 2: Comparison of SEIFC airflow measurements from aerobic and anoxic zones with blower data

We employed blower data sourced from the plant supervisory control and data acquisition (SCADA) system alongside airflow measurements. The blower data was used to evaluate the $N_2O$ emission rates from the nitrifying system; however, the absence of a blower in the anoxic zone eliminates the need to quantify emissions and creates a significant knowledge gap in addressing the overall emission rate from the activated sludge tank (AST) and emission fraction (EF) from the WWTPs. **Figure S1** shows that when considering blower data to quantify $N_2O$ emission from the aerobic zone, the emissions were overestimated and failed to identify the anoxic zone when compared to the airflow meter measurements. Utilizing an airflow meter with our established protocol has facilitated the measurement of $N_2O$ concentration and flux from both aerobic and anoxic zones with ease[1,2].

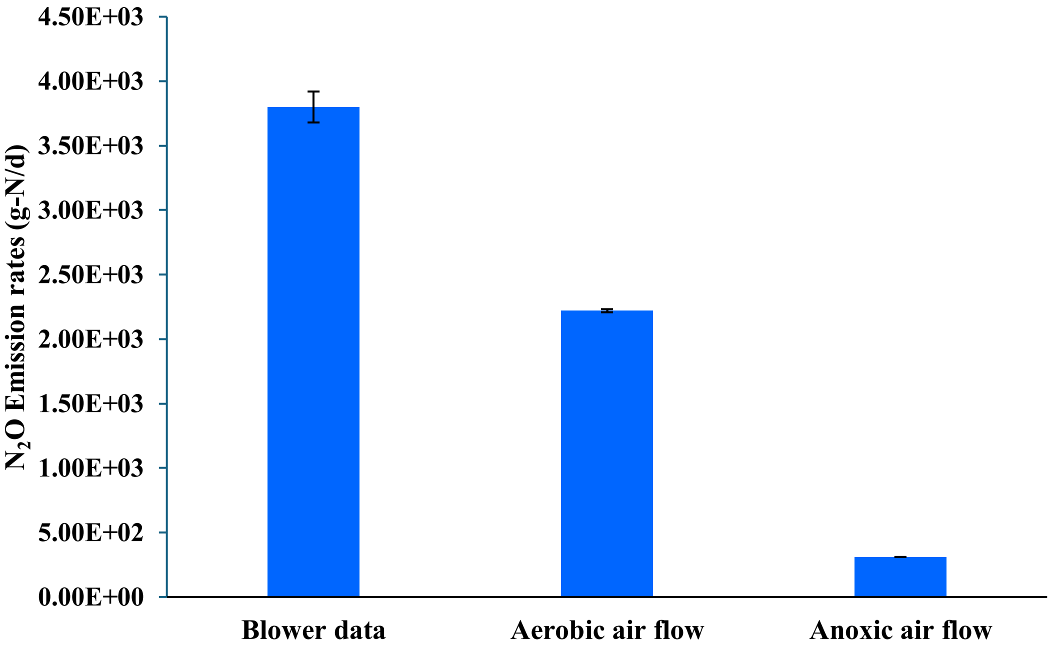


**Figure S1.** The plot represents the $N_2O$ emission rates quantified using blower and airflow data measured from the aerobic and anoxic zones in a full-scale activated sludge tank.

This study also quantified the off-gas flow rates (L $min^{-1}$) from different aerobic zones in the step-feed BNR configuration **(Figure S2).** Blower off-gas flow rates were higher than directly measured off-gas flow rates. The results show that considering only blower off-gas flow rates can overestimate the emission rates when compared to airflow data from the flux chamber, leading to inaccurate estimation of $N_2O$ emission rates and EF measurements.

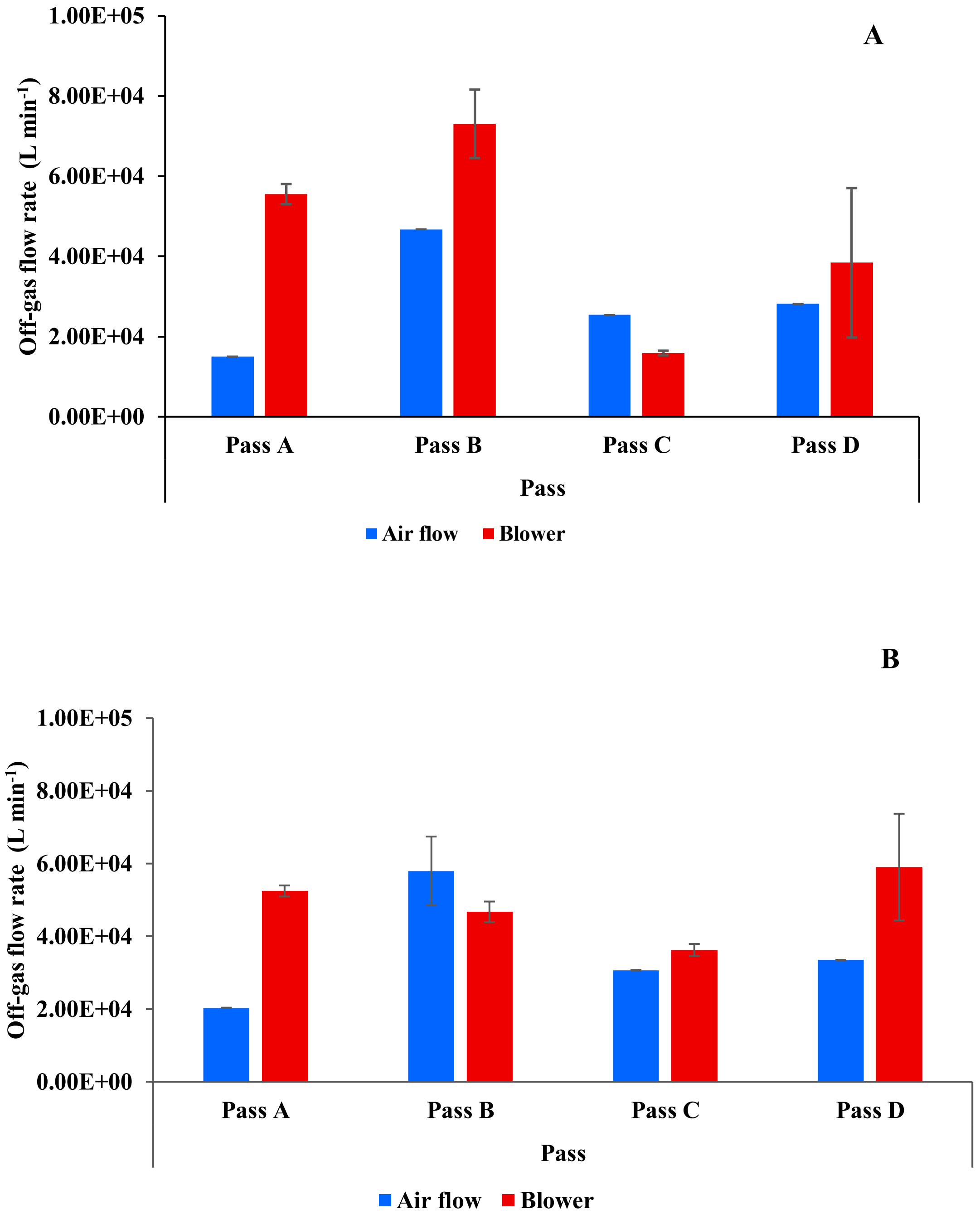


**Figure S2.** The plot represents the off-gas flow rate in the aerobic zone using air blower data (scfm) *vs* airflow meter (m/s). **(A)** represents the off-gas flow rate calculated during the winter campaign (18±3°C), **(B)** represents the off-gas flow rate calculated during the summer campaign (23±3°C).

Furthermore, different sweep-airflow rates were introduced into the SEIFC in the anoxic zone to measure the off-gas flow rate (**Figure S3**). The suitability of employing a sweep-air flow rate of 5 L $min^{-1}$ was confirmed based on comparison with the measured off-gas flow rate.

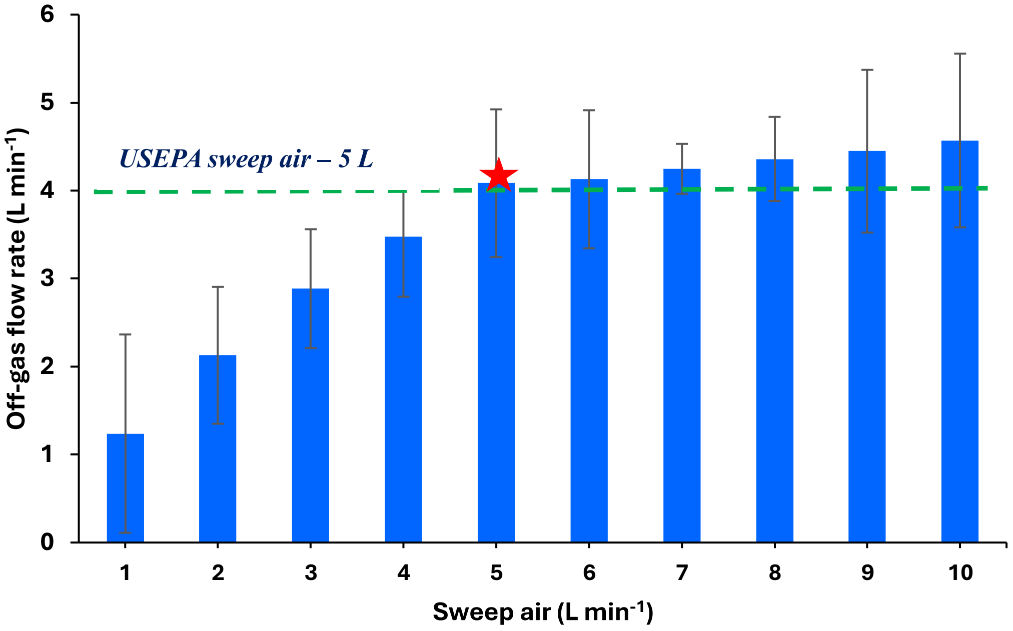


**Figure S3.** The plot represents the off-gas flow rate (L $min^{-1}$) at different sweep-airflow rates (L $min^{-1}$) introductions to the anoxic zone flux chamber setup.

**Section 3: Hyperparameter tuning for predicting $N_2O$ flux using different ML models**

**Table S2. Hyperparameters for ML models tested**

| Random Forest – Aerobic | | | |
|---|---|---|---|
| **Parameters** | **Minimum** | **Maximum** | **Selected** |
| Number of predictors at each split | 2 | 5 | 5 |
| Number of estimators (trees) | 100 | 800 | 500 |
| Minimum node size | 1 | 10 | 10 |
| Max depth of base trees | 10 | 40 | 30 |
| **Random Forest: Anoxic** | | | |
| **Parameters** | **Minimum** | **Maximum** | **Selected** |
| Number of predictors at each split | 2 | 5 | 4 |
| Number of estimators (trees) | 100 | 800 | 500 |
| Minimum node size | 1 | 10 | 5 |
| Max depth of base trees | 10 | 40 | 40 |
| **Gradient Boosting: Aerobic** | | | |
| **Parameters** | **Minimum** | **Maximum** | **Selected** |
| Number of estimators (trees) | 50 | 200 | 200 |
| Learning rate | 0.01 | 1 | 0.1 |
| Max depth of base trees | 3 | 9 | 7 |
| **Gradient Boosting: Anoxic** | | | |
| **Parameters** | **Minimum** | **Maximum** | **Selected** |
| Number of estimators (trees) | 50 | 200 | 200 |
| Learning rate | 0.01 | 1 | 0.1 |
| Max depth of base trees | 3 | 9 | 5 |

| Extreme Gradient Boosting Model (XGBoost): Aerobic | | | |
|---|---|---|---|
| **Parameters** | **Minimum** | **Maximum** | **Selected** |
| Number of estimators (trees) | 50 | 1200 | 400 |
| Learning rate | 0.01 | 1 | 0.1 |
| Minimum child weight | 1 | 5 | 4 |
| Subsample | 0.2 | 0.5 | 0.2 |
| Max depth of trees | 1 | 9 | 7 |
| Splitting criterion | Squared error | | Squared error |

| Extreme Gradient Boosting Model (XGBoost): Anoxic | | | |
|---|---|---|---|
| **Parameters** | **Minimum** | **Maximum** | **Selected** |
| Number of estimators (trees) | 50 | 1200 | 800 |
| Learning rate | 0.01 | 1 | 0.1 |
| Minimum child weight | 1 | 5 | 1 |
| Subsample | 0.2 | 0.8 | 0.8 |
| Max depth of trees | 1 | 7 | 5 |
| Splitting criterion | Squared error | | Square error |

### Section 4: Model performance under different machine learning models

A summary of the predictive performance of the optimized models is presented below. Model performance for predicting $N_2O$ flux, emission rates, and concentration was evaluated on the held-out test dataset using $R^2$, MSE, RMSE and MAE.

**Table S3. Performance evaluation of ML models for aerobic and anoxic zones**

**a. Aerobic zone ML model evaluation:**

| Model | Dependent variables | Performance metrics (Held-out test) | | | |
|---|---|---|---|---|---|
| | | $R^2$ | MSE | RMSE | MAE |
| Random Forest | $N_2O$ flux (g-N/m$^2$) | 0.88 | 2.56 | 1.60 | 0.50 |
| | $N_2O$ Emission rate (g-N/d) | 0.37 | 5042954 | 2245.652 | 750.91 |
| | $N_2O$ concentration (ppm) | 0.72 | 76.0 | 8.71 | 2.50 |
| Gradient Boosting | $N_2O$ flux (g-N/m$^2$) | 0.97 | 0.53 | 0.72 | 0.24 |
| | $N_2O$ Emission rate (g-N/d) | 0.84 | 1226446 | 1107.45 | 273.05 |
| | $N_2O$ concentration (ppm) | 0.78 | 60.6 | 7.78 | 2.01 |
| XGBoost | $N_2O$ flux (g-N/m$^2$) | 0.97 | 0.48 | 0.69 | 0.18 |
| | $N_2O$ Emission rate (g-N/d) | 0.89 | 840286.1 | 916.67 | 159.26 |
| | $N_2O$ concentration (ppm) | 0.71 | 80.65 | 8.90 | 2.0 |

**b. Anoxic zone ML model evaluation:**

| Model | Dependent variables | Performance metrics (Held-out test) | | | |
|---|---|---|---|---|---|
| | | $R^2$ | MSE | RMSE | MAE |
| Random Forest | $N_2O$ flux (g-N/m$^2$) | 0.75 | 0.07 | 0.26 | 0.04 |
| | $N_2O$ Emission rate (g-N/d) | 0.75 | 4142 | 64.36 | 10.96 |
| | $N_2O$ concentration (ppm) | 0.96 | 0.48 | 0.69 | 0.24 |
| Gradient Boosting | $N_2O$ flux (g-N/m$^2$) | 0.75 | 0.07 | 0.27 | 0.04 |
| | $N_2O$ Emission rate (g-N/d) | 0.97 | 4125 | 20.31 | 4.76 |
| | $N_2O$ concentration (ppm) | 0.98 | 0.21 | 0.46 | 0.179 |
| XGBoost | $N_2O$ flux (g-N/m$^2$) | 0.87 | 0.04 | 0.20 | 0.03 |
| | $N_2O$ Emission rate (g-N/d) | 0.95 | 708.5 | 26.61 | 5.63 |
| | $N_2O$ concentration (ppm) | 0.98 | 0.16 | 0.41 | 0.15 |

**Section 5: Relationship between % Nitrogen removed and $N_2O$ EF% for mainstream and sidestream process configurations**

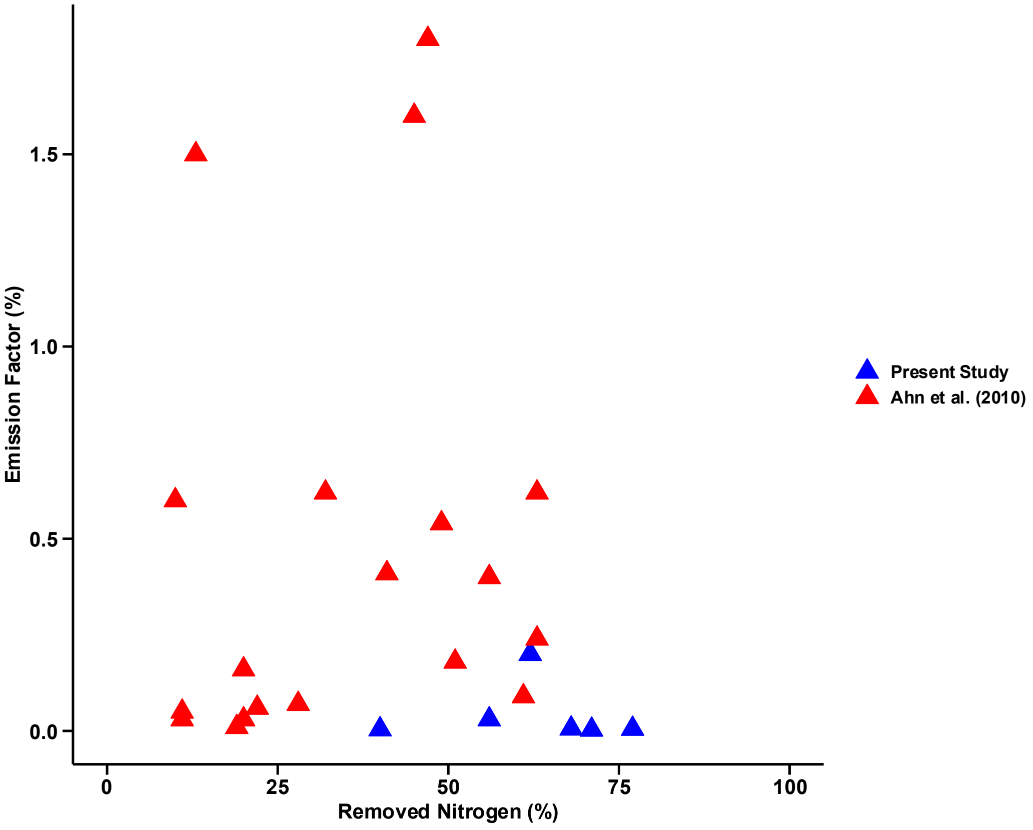


**Figure S4.** Relationship between % Nitrogen removed and Emission Factor (EF)% for mainstream and sidestream systems from the present study (▲blue triangle) and previous study[1] (▲red triangle). *The sidestream SHARON EF and % removed nitrogen from the present study was excluded from the plot because of its considerably higher EF compared to mainstream configurations.*

### Section 6: Pearson correlation between $N_2O$ flux and process operational variables

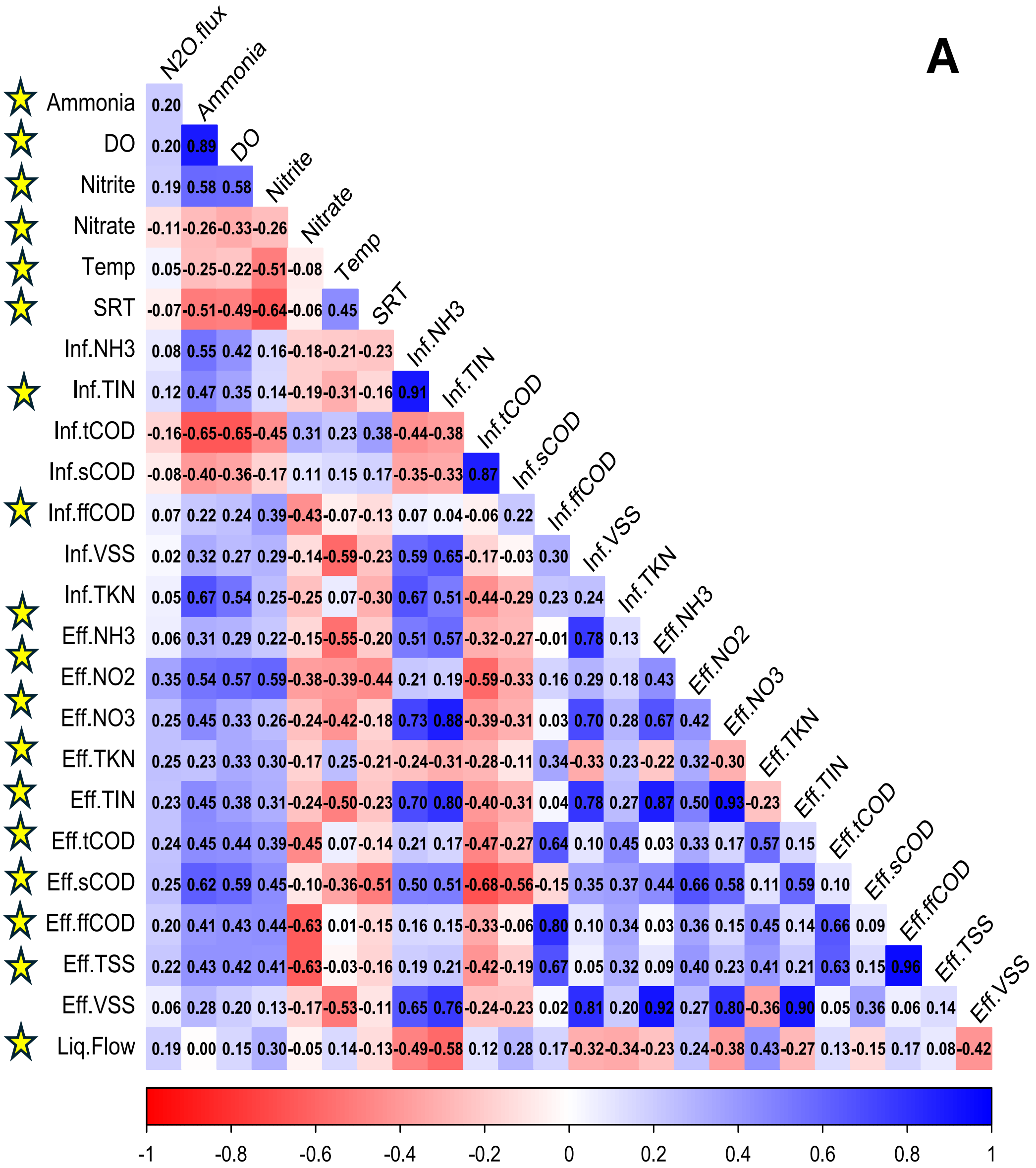

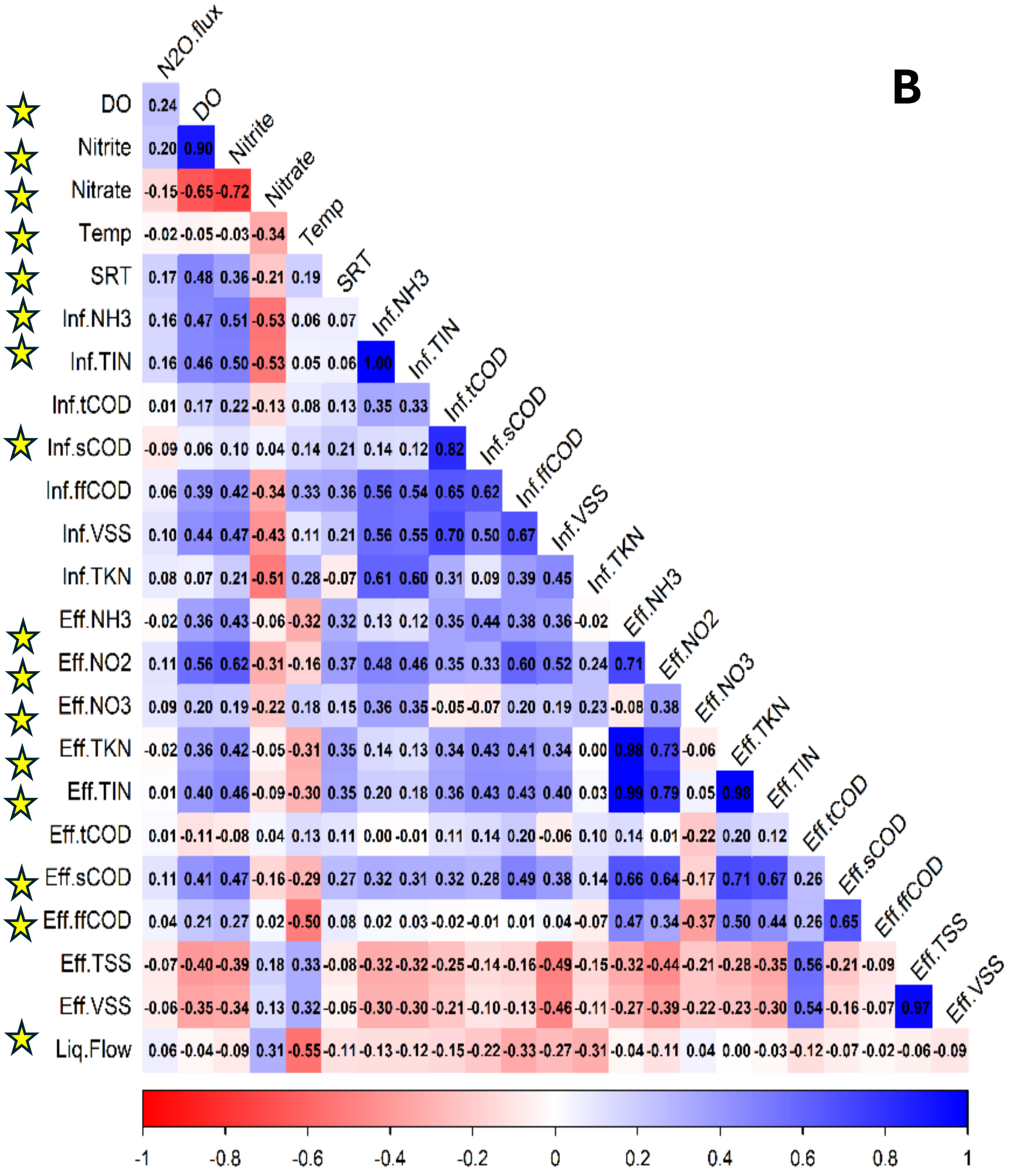


**Figure S5.** Pearson correlation coefficients between $N_2O$ flux and process operational variables for the aerobic **(A)** and anoxic **(B)** zones. Stars indicate variables with statistically significant correlations with $N_2O$ flux ($p<0.05$).

### Section 7: Permutation importance from XGBoost model on $N_2O$ fluxes for both aerobic and anoxic zones

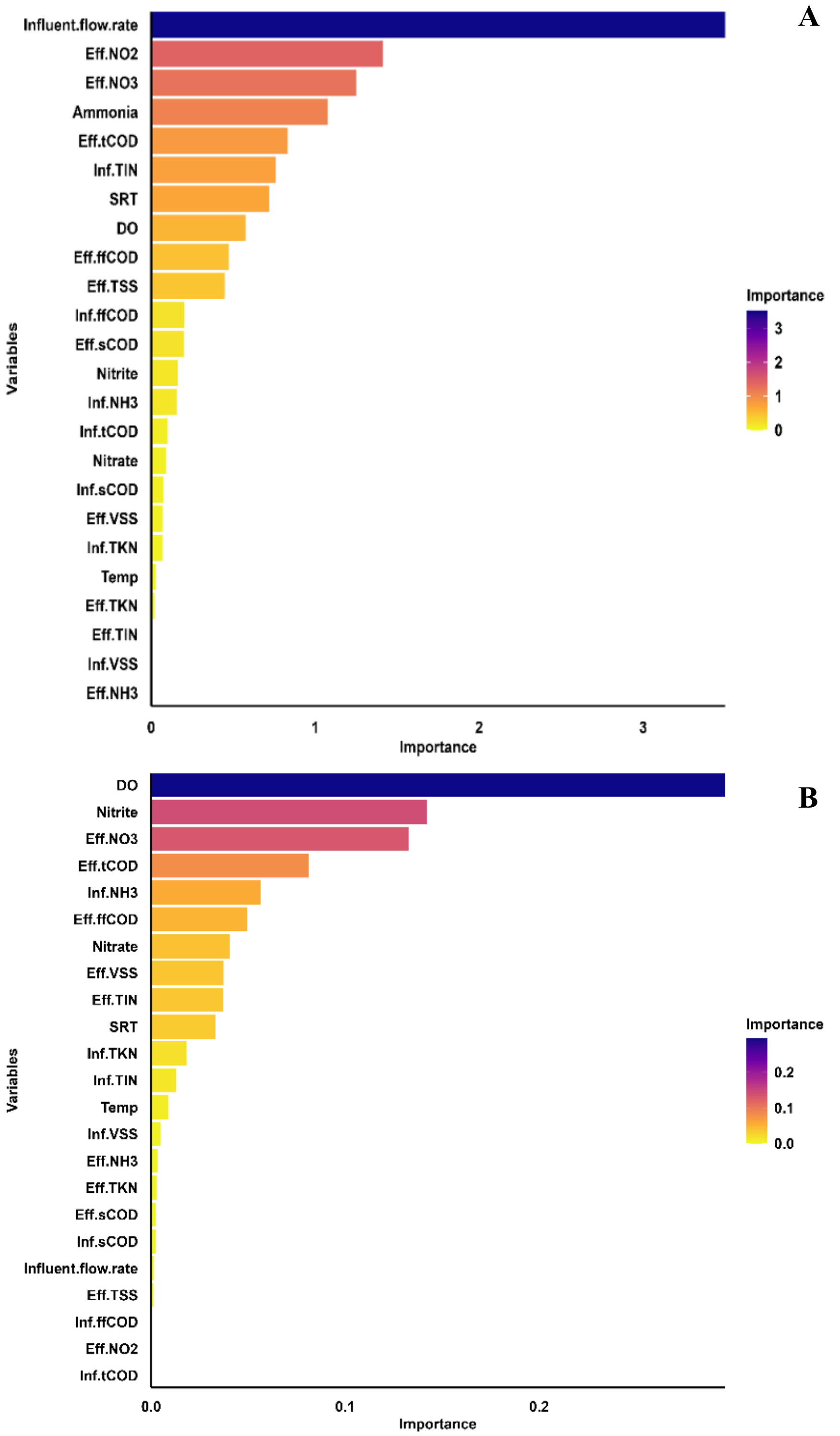


**Figure S6.** Permutation importance on $N_2O$ flux trained on XGBoost model for both **(A)** aerobic (24 variables) and **(B)** anoxic zones (23 variables).

### Section 8: SHAP analysis from XGBoost model on $N_2O$ emission rates and concentrations for aerobic and anoxic zones

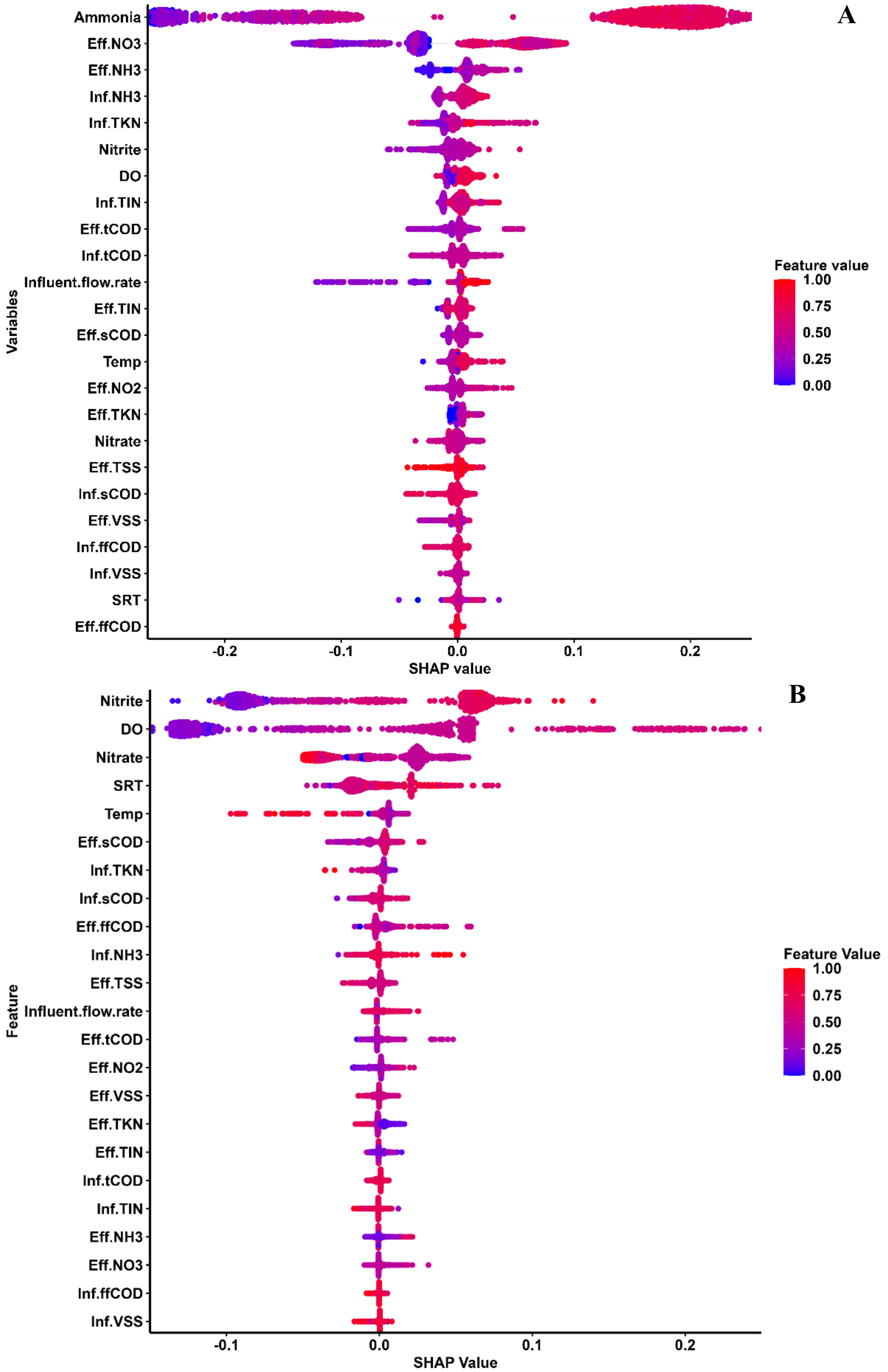

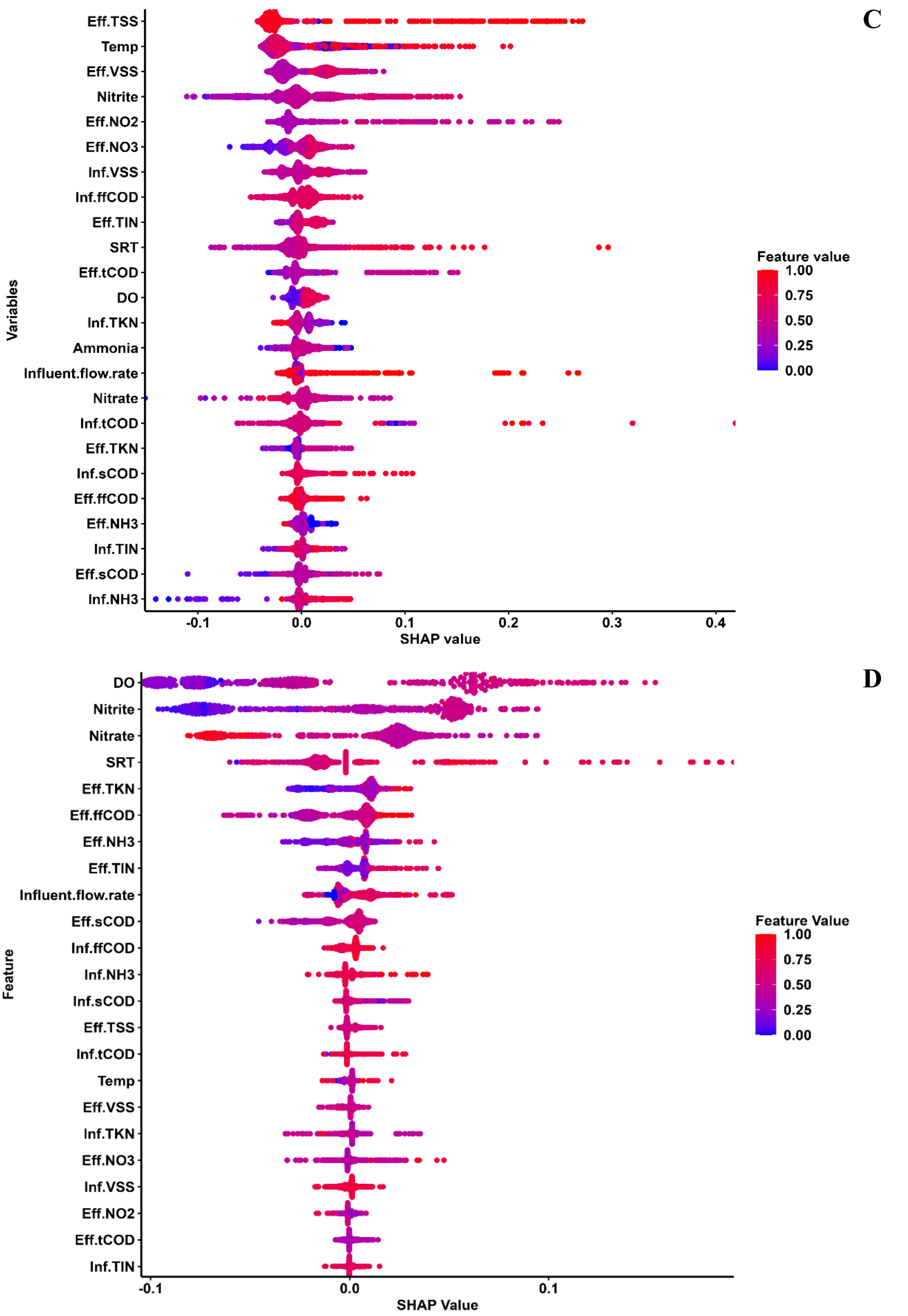


**Figure S7.** Beeswarm SHAP analysis on $N_2O$ emission rate **(A, B)** and $N_2O$ concentration **(C,D)** trained on XGBoost model for both aerobic **(A, C)** and anoxic **(B, D)** zone

**Table S4. Predicted $N_2O$ flux ratios from the mechanistic consistency assessment using paired-variable scenario analyses in aerobic and anoxic zones.**

| **Zone** | **Interaction** | **High Emission scenario** | **Low emission scenario** | **Median ratio** | **Lower 95% CI** | **Upper 95% CI** |
|---|---|---|---|---|---|---|
| **Aerobic** | SRT x $NH_3$ | Short SRT + High $NH_3$ | Long SRT + Low $NH_3$ | 8.91 | 3.22 | 38.72 |
| | DO x $NH_3$ | Low DO + High $NH_3$ | High DO + Low $NH_3$ | 7.83 | 2.22 | 52.82 |
| **Anoxic** | DO x $NO_2^-$ | High DO + High $NO_2^-$ | Low DO + Low $NO_2^-$ | 8.80 | 3.83 | 21.33 |
| | DO x SRT | High DO + Short SRT | Low DO + Long SRT | 5.15 | 1.47 | 18.80 |

### Process schematics of WWTPs reactors sampled

### STEP-FEED BNR

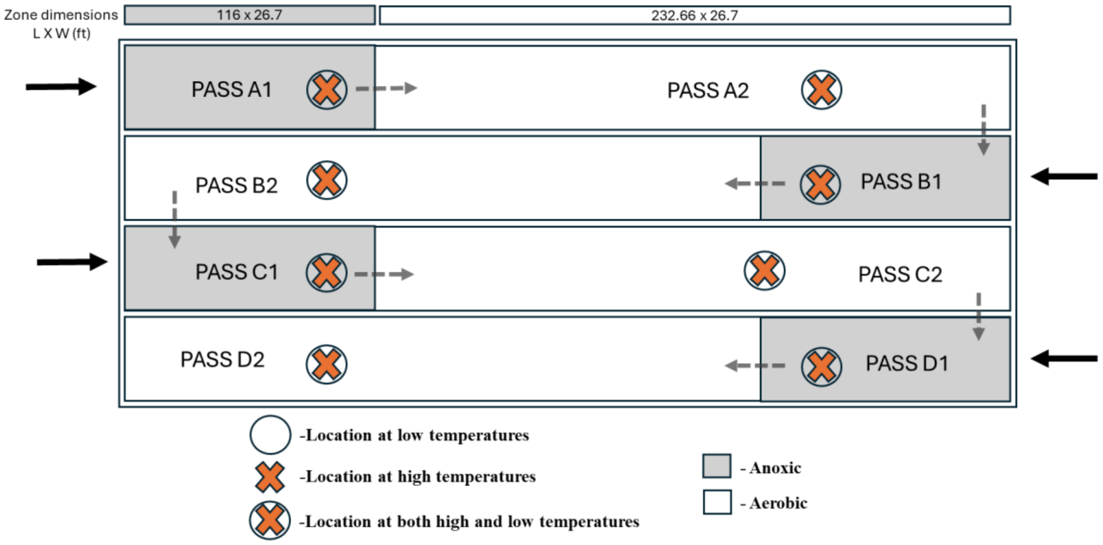


### AMMONIA BASED AERATION CONTROL (ABAC)

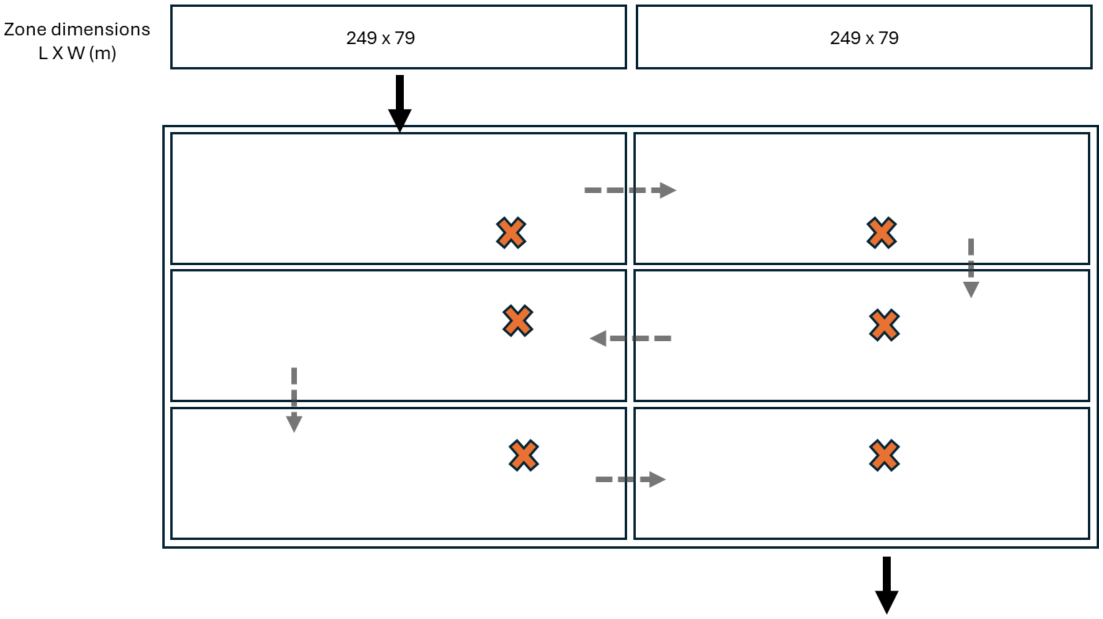

### PARTIAL DENITRIFICATION ANAMMOX (PDNA)

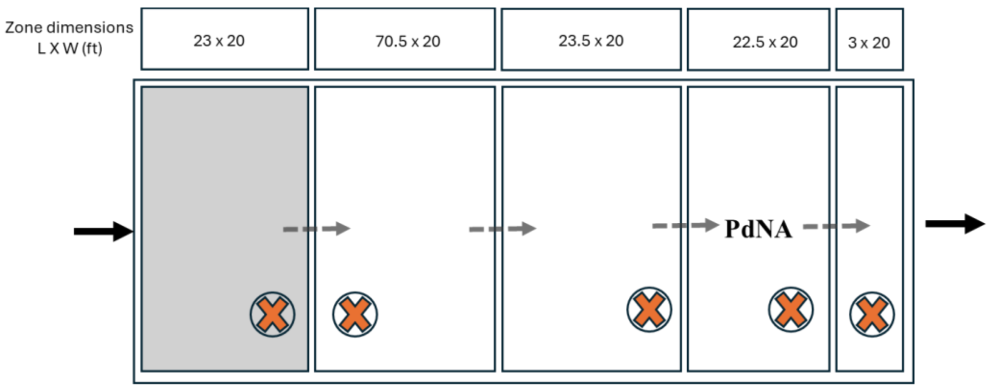


### SHARON

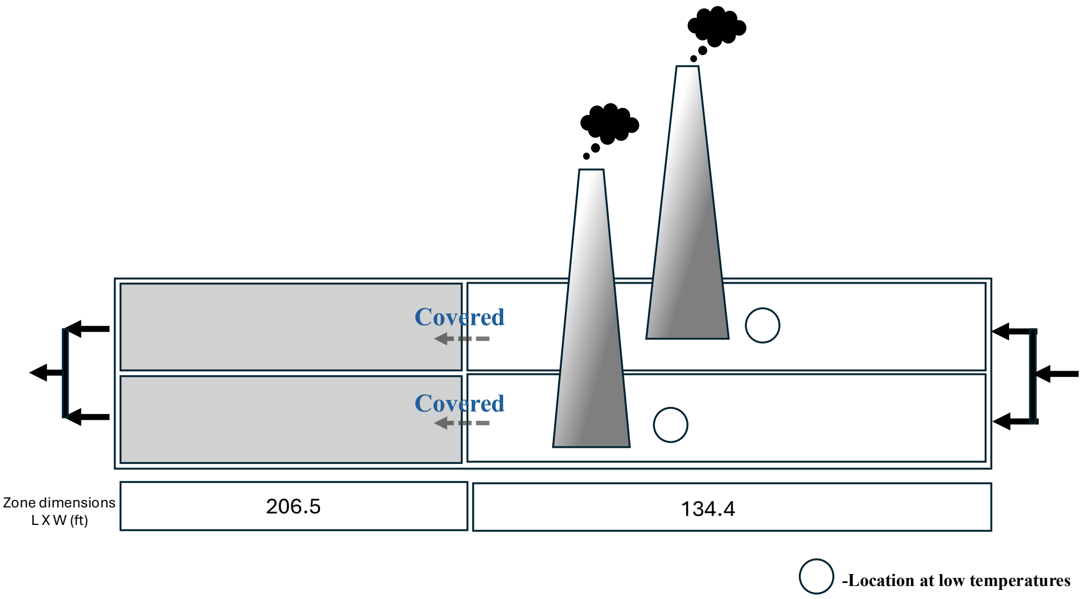

The preprocessing pipeline was designed to prevent information leakage. Following the train–test split, *missForest* was fit exclusively on the training predictors. The learned imputations were applied only to the training data, whereas missing values in the independent test set were imputed using summary statistics (medians) derived solely from the training data. The relevant code snippet is shown below:

```
library(missForest)

# missForest AFTER split
# Only impute predictor columns in training
train_predictors <- train_raw[, pred_cols]

set.seed(123)
mf_train <- missForest(
  train_predictors,
  maxiter = 10,
  ntree = 500,
  verbose = TRUE
)

# Replace training predictors with imputed values
train_raw[, pred_cols] <- mf_train$ximp

# Impute TEST using training information
train_medians <- sapply(
  train_raw[, pred_cols],
  median,
  na.rm = TRUE
)

for (v in pred_cols) {
  test_raw[[v]][is.na(test_raw[[v]])] <- train_medians[v]
}


# 4. Skewness detection on TRAIN only

skew_vals <- sapply(train_raw[, pred_cols, drop = FALSE], function(x) {
  e1071::skewness(x, na.rm = TRUE, type = 2)
})

vars_to_log <- names(skew_vals[is.finite(skew_vals) & abs(skew_vals) > 1])

print("Variables log-transformed based on train-set skewness:")
print(vars_to_log)
```

### REFERENCES


1. Ahn, J. H. *et al.* N2O emissions from activated sludge processes, 2008-2009: Results of a national monitoring survey in the united states. *Environmental Science and Technology* **44**, 4505–4511 (2010).
2. Chandran, K. *Protocol for the Measurement of Nitrous Oxide Fluxes from Biological Wastewater Treatment Plants*. *Methods in Enzymology* vol. 486 (Elsevier Inc., 2011).
3. Andrew D. Eaton, American Water Works Association, Water Environment Federation. *Standard Methods for the Examination of Water and Wastewater*. (APHA-AWWA-WEF, Washington, D.C., 2005).
